# Community context supersedes species identity in bacterial interactions and disease outcomes in the tomato phyllosphere

**DOI:** 10.64898/2026.01.16.699983

**Authors:** Tiffany N. Batarseh, Jose O. Collado, Chuyu Wang, Elijah C. Mehlferber, Rocelia M. Alvarez-Navarrete, Fiona J. Wagner, Britt Koskella

## Abstract

Microbial community function is challenging to predict from the properties of individual members alone as ecological interactions can alter taxon behavior. Using a 16-member synthetic bacterial community from the tomato phyllosphere, we ask whether interaction outcomes at the community level can be predicted based on pairwise interactions by combining whole-genome sequencing, phenotypic profiling, and reciprocal spent-media assays spanning over 600 single-species and community-level combinations. Community-conditioned media reduced growth substantially more than the average of the member strains effects, but never more than the most suppressive member. The number of members failed to predict suppression, indicating that composition rather than richness drove these effects. Neither genomic, growth, nor phylogenetic traits predicted interaction outcomes, even in cross-validated models. In vivo, no single strain significantly reduced disease caused by the pathogen, *Pseudomonas syringae* on tomato seedlings, but microbial communities were disease suppressive. This was in stark contrast to our in vitro results, where the pathogen behaved like a commensal, most suppressed by the same single strains that also suppressed the other members (notably *E. sibiricum* SC13) and was no more sensitive to community filtrates than the commensals were, indicating that this protection was not reducible to the chemical environment the community generates. Overall, we find that interaction outcomes are governed by the composition of the community, competitive outcomes, and are hard to predict based on phenotypic or genomic properties of the individual taxa, indicating that the community, in the context of its host, should be considered when predicting microbiome function.

## Introduction

The function of a microbial community is shaped by the taxonomic and metabolic diversity of its members as well as the interactions among them. Microbial interactions such as competition for a shared limiting resource, secretion of antimicrobial toxins like antibiotics/bacteriocins, or chemical modification of the environment determine not only whether taxa coexist, but also whether communities are more or less likely to resist invasion, how stable host-associated microbiomes are over time, and which collective functions might emerge [1–3]. Because these interactions are highly context dependent and can act through multiple mechanisms, community-level outcomes often cannot be inferred from either taxonomic composition or knowledge about pairwise interactions alone [4, 5].

Although individual microbial traits, such as growth rate, carbon usage, environmental tolerance, or biosynthetic potential are encoded in microbial genomes, linking these traits to phenotypes within a given environment and/or to interaction outcomes remains difficult [6–9]. Genomic relatedness, especially at broad scales such as 16S rRNA gene similarity, does not necessarily translate into similar ecological behavior or interaction phenotypes [10]. Comparative genomics approaches have successfully linked genome size, metabolic breadth, and functional gene content to broad ecological strategies [11, 12], yet whether such additive summaries capture interaction outcomes has rarely been tested. Moreover, interactions observed in isolation often fail to predict behavior in more complex community contexts [9], highlighting the importance of indirect and higher-order effects. The challenge of prediction is especially apparent in disease-protective microbiomes.

Pathogen suppression has traditionally been studied either at the level of single biocontrol agents or through correlations with overall community diversity, leaving a gap in understanding how interactions among intermediate subsets of community members generate protective outcomes [13, 14]. Neither monoculture nor whole-community assays can separate a community property from a member property, and pairwise measurements rarely resolve it either, since the function of an assembled consortium frequently departs from what its constituent pairs predict [15]. Removing single members from an otherwise intact community can show whether a given strain is necessary for protection. In the human gut, dissecting protection across defined community subsets has shown that communities restrict pathogen growth far more than their individual members and that the degree of nutrient-use overlap between residents and invader predicts the outcome [16]. Whether the same structure holds in the other microbiomes such as the plant phyllosphere, where nutrient availability is patchier and colonization is more stochastic, has not been tested. Protection that exceeds what members achieve alone could arise from collective resource use that leaves little niche space for an invader, from direct antagonism by particular strains, or from modification of the shared chemical environment, for example through pH or inhibitory metabolites.

Spent media assays, where bacteria are grown for a finite period and then live cells are removed through filtration, provide a tractable experimental framework for quantifying environmentally mediated interactions among bacteria by measuring how prior community membership alters growth conditions for subsequent colonizers. Because most colonizers arrive in an environment that already hosts other taxa, their establishment depends largely on which resources remain and what metabolites, toxins, or phages have accumulated in the local environment [17]. These assays have shown great promise in describing the *in vitro* interaction networks of the mouse gut, identifying keystone species driving community outcomes, and predicting *in vitro* assembly of a 15-member community [18, 19]. However, they are typically run as pairwise ‘invasions’ and may miss community-level effects, because bacteria grown together can modify the chemical environment in ways not predictable from their individual spent media.

Here, we use PhylloStart, a defined synthetic community of tomato phyllosphere bacteria (Supplementary Table 1; [20]). Synthetic communities (SynComs) of cultured, well-described isolates provide a reductionist tool for dissecting pairwise and community-level interactions in a controlled and tractable framework [21]. PhylloStart contains most genera previously identified as core tomato microbiome members [22] and maintains important functions, including improving tomato reproductive success and protecting tomato against the model pathogen *Pseudomonas syringae*, although this protection is conditional [20, 23]. Combining reciprocal spent-media growth assays, phenotypic profiling and genomics of the entire SynCom, we test the extent to which microbial ‘invasion’ can be predicted by the genomic features of the bacteria, their metabolic overlap, and/or higher-order community interactions. If protection against *P. syringae* is intrinsic to how these bacteria interact, the pathogen should be distinctively sensitive to community-conditioned media; if it requires the leaf, it should be indistinguishable from the commensals in a cell-free assay. We therefore treated *P. syringae* as one focal strain among seventeen and tested the same communities on seedlings. By connecting pairwise and community-scale outcomes with the genotypic and phenotypic information of each member, this work contributes to efforts to predict and engineer microbiome function.

## Results

### Genomic and phenotypic characterization of the phyllosphere bacteria

The 16 commensals of PhylloStart, isolated from tomato plants to represent the breadth of tomato phyllosphere diversity [20, 22], span 11 genera in three phyla: eight Pseudomonadota (Gammaproteobacteria, except the betaproteobacterium *Massilia aurea* SC09), five Actinomycetota and three Bacillota (Supplementary Table 1). Hybrid genome assemblies ranged from 3.1 to 6.6 Mb (mean of 4 contigs) and from 34% to 73% GC (Supplementary Table 2), with genome completeness and contamination assessed by CheckM (Supplementary Table 3). Phylogenies from the 16S rRNA gene and from 107 conserved single-copy genes were largely congruent, and pairwise average nucleotide identity reflected this structure (Figure 1A; Supplementary Figures 1–2; Supplementary Results). BlastKOALA annotated 53.7% of genes per genome on average (range 39.7–67.3%). Only about a tenth of the gene content of an average genome was shared by all strains, so any predictive signal should come from where strains differ rather than from what they share (Supplementary Results). Predicted amino acid biosynthesis and small-carbon-source catabolism also varied among strains, with greater diversity in catabolic potential (Supplementary Figures 3–4). Genomes encoded 55 to 147 carbohydrate-active enzymes (CAZymes), a count unrelated to genome size (Figure 1A; Supplementary Results). Because iron competition operates without cell contact and siderophores persist in cell-free filtrates, iron-acquisition capacity is a candidate predictor of the interactions measured here; *Pseudomonas* genomes carried the most iron-related genes (129 on average, against 61 for all others; Figure 1A). These counts index annotated capacity rather than realized function: about half of the coding capacity of these genomes is unannotated, and carrying iron-acquisition genes does not imply siderophore production under a given condition. Secretion systems were most common in the Gram-negative strains, and 138 secondary metabolite biosynthetic gene clusters were identified across the 17 genomes, from 3 in *E. sibiricum* SC13 to 13 in *B. thuringiensis* SC35 (**Supplementary Figure 5; Supplementary Results**).

**Fig 1.**
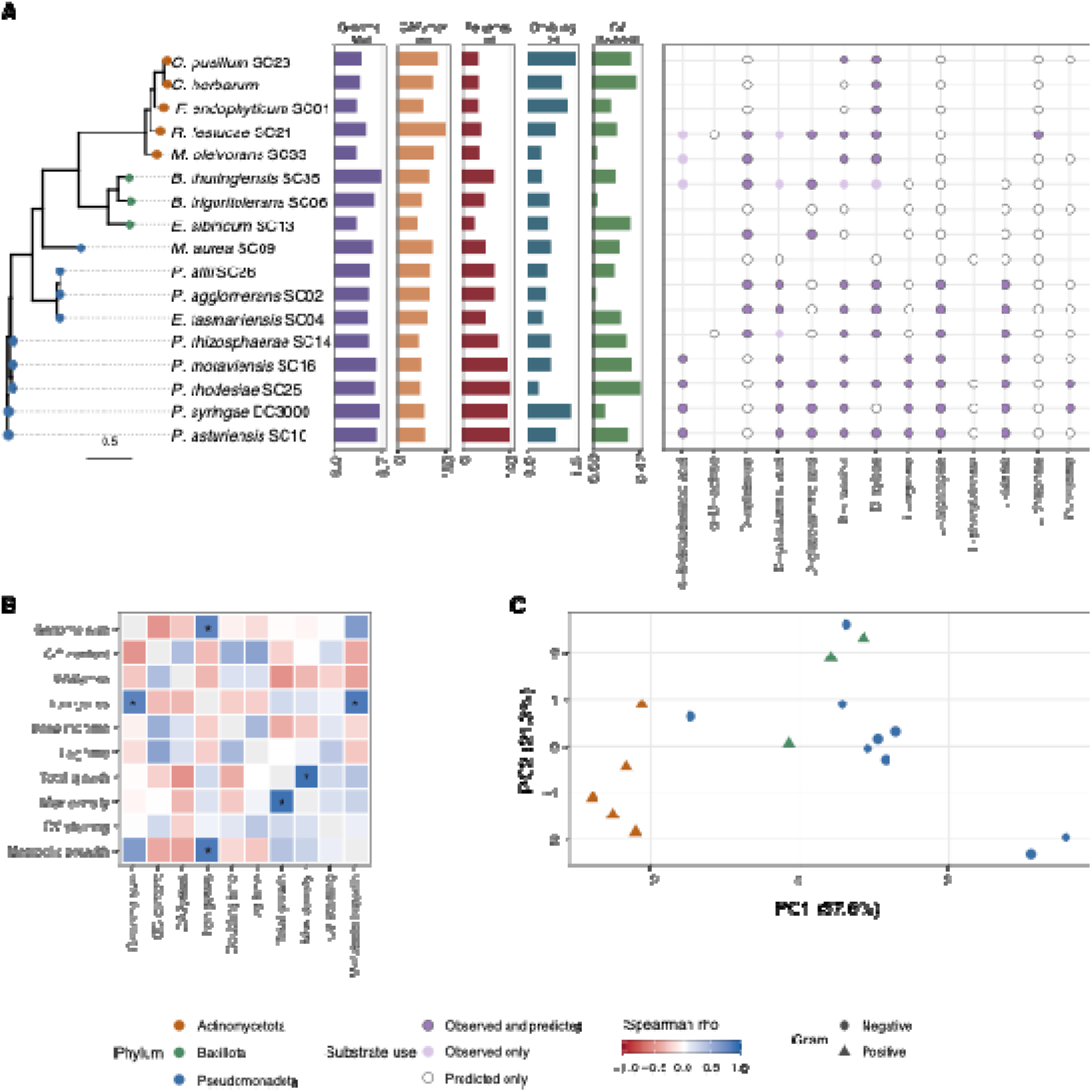
Genomic and phenotypic characterization of the 17 phyllosphere isolates in phylogenetic context. (A) Core-gene phylogeny of all 17 isolates, with tips colored by phylum, and aligned tracks giving genome size, CAZyme count, iron-related gene count (FeGenie), doubling time in KB, biofilm formation, and carbon or amino acid source use. Bar color indicates which trait is plotted; bar length is the value of that trait, on an independent scale per trait. In the substrate panel a filled circle indicates use observed on Biolog EcoPlates, an open circle indicates use predicted from the genome annotation but not observed, and substrates are restricted to the 13 for which a genomic prediction was available. Genus names are abbreviated; full names and taxonomic assignments are given in Supplementary Table 1. (B) Spearman correlation matrix among the ten continuous traits. Cell values are correlation coefficients; asterisks mark pairs significant at q < 0.05 after Benjamini-Hochberg correction across all unique pairs, and unmarked cells are not significant. (C) Principal components analysis of all standardized traits, with points colored by phylum and shaped by Gram status. Full amino acid biosynthesis and small carbon utilization predictions are given in Supplementary Figures 3–4, and the relationship between CAZyme count and genome size is reported in the Supplementary Results.

Because life-history traits are predicted to influence community assembly and function [8], we also measured growth, biofilm formation and carbon-source use. Growth in King’s B medium varied widely, and all strains reached stationary phase within 72 h (Table 1; Supplementary Figure 6); 11 of 17 strains were fast growers (mean doubling time < 0.75 h) and 6 were slow growers (> 0.75 h). Eleven of the 17 strains formed biofilms (Table 1; Figure 1A). On Biolog EcoPlates, carbon-source use clustered by phylogeny (Figure 1A; Supplementary Figure 7) and ranged from 9 of 31 substrates (*F. endophyticum* SC01) to 24 of 31 (*M. oleivorans* SC33). Several traits covaried across strains: iron-related gene content rose with both genome size and metabolic breadth, and total growth with maximum density (Figure 1B), and strains grouped largely by phylum in a principal components analysis of all traits (Figure 1C).

**Table 1.** Growth rate characteristics of the 17 phyllosphere associated bacteria.

| **Species** | **Isolate** | **Mean AUC\*** | **Mean Lag Time** | **Mean Doubling Time** | **Mean Maximum Density** | **Biofilm Formation<sup>1</sup>** |
|---|---|---|---|---|---|---|
| *Pseudomonas syringae* | DC3000 | 18.4 | 0.58 | 1.35 | 0.407 | N |
| *Frigoribacterium endophyticum* | SC01 | 40.4 | 9.34 | 1.25 | 1.15 | N |
| *Pantoea agglomerans* | SC02 | 56.8 | 3.27 | 0.549 | 1.42 | N |
| *Curtobacterium herbarum* | SC03 | 24.1 | 7.75 | 1.08 | 0.708 | Y |
| *Erwinia tasmaniensis* | SC04 | 38.5 | 1.18 | 0.458 | 0.947 | Y |
| *Brevibacterium*<sup>2</sup> *frigoritolerans* | SC06 | 34 | 4.94 | 0.651 | 0.873 | N |
| *Massilia*<sup>3</sup> *aurea* | SC09 | 20.1 | 6.96 | 0.698 | 0.531 | Y |
| *Pseudomonas asturiensis* | SC10 | 34.1 | 8.6 | 0.902 | 0.748 | Y |
| *Exiguobacterium sibiricum*<sup>4</sup> | SC13 | 41.7 | 3.4 | 0.608 | 0.997 | Y |
| *Pseudomonas rhizosphaerae* | SC14 | 40.3 | 2.92 | 0.69 | 1.05 | Y |
| *Pseudomonas moraviensis* | SC16 | 73.1 | 7.41 | 0.697 | 1.71 | Y |
| *Rathayibacter festucae* | SC21 | 26.4 | 6.27 | 0.893 | 0.997 | N |
| *Curtobacterium pusillum* | SC23 | 36 | 6.21 | 1.51 | 1 | Y |
| *Pseudomonas rhodesiae* | SC25 | 73.1 | 5.24 | 0.333 | 1.62 | Y |
| *Pantoea allii* | SC26 | 51.1 | 3.43 | 0.59 | 1.18 | Y |
| *Microbacterium oleivorans* | SC33 | 26.7 | 4.18 | 0.409 | 0.733 | N |
| *Bacillus thuringiensis* | SC35 | 30.3 | 3.03 | 0.431 | 0.861 | Y |

\*Area under the curve; measured to represent the total bacterial growth
<sup>1</sup>Biofilm formation (Y = yes and N = no) as identified by the crystal violet test and a statistically significant difference from the control in crystal violet staining
<sup>2</sup>Synonymous with *Peribacillus*
<sup>3</sup>Synonymous with *Telluria*
<sup>4</sup>Synonymous with *Exiguobacterium artemiae*

### Interaction effects in spent media

We grew all 17 strains in 17 single-strain and 23 community spent media (filtrates; Supplementary Table 4). The community panel comprised the intact 16-member PhylloStart community, the same community plus *P. syringae*, sixteen single-strain drop-outs, and five group drop-outs removing all *Pseudomonas*, all Gram-negative members, all Gram-positive members, both *Pantoea*, or two of the four *Pseudomonas* (*P. moraviensis* SC16 and *P. rhodesiae* SC25). Growth in each filtrate was expressed relative to growth in fresh medium (d_AUC_; Methods).

Filtrates reduced growth on average (mean d_AUC_ =-0.29; Supplementary Figure 8), community filtrates more than single-strain filtrates (-0.35 versus-0.14; Figure 2A). In a linear mixed-effects model (Methods), community filtrates reduced growth by 0.203 d_AUC_ units more than single-strain filtrates (df = 38.5, P = 2.2 × 10□□; Supplementary Figure 9). A conservative, model-free check agreed: comparing each focal strain’s mean d_AUC_ in community and in single-strain filtrates, community filtrates were more suppressive for 16 of 17 focal strains (median difference =-0.115, Wilcoxon signed-rank P = 3.1 × 10□□). Plate effects did not explain this difference (Supplementary Figure 10; Supplementary Results).

**Fig 2.**
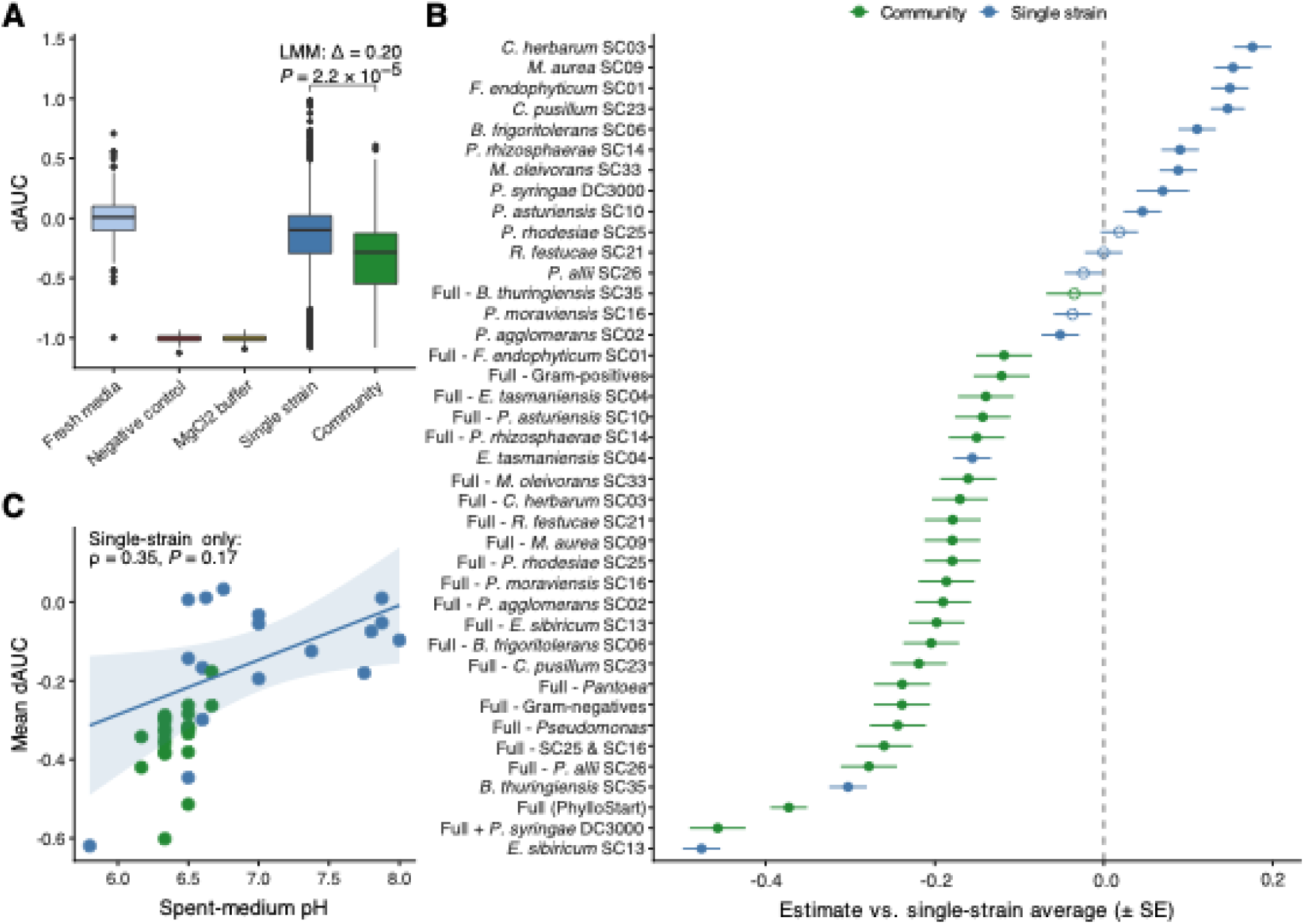
Community filtrates suppress growth more than single-strain filtrates, and suppression is not explained by pH. Relative growth change (d_AUC_) is each focal strain’s growth in a spent medium relative to its own growth in fresh King’s B medium; zero indicates growth matching fresh medium and negative values indicate suppression. A) d_AUC_ by medium class across all focal strains, for fresh medium, the MgClO buffer and negative controls, the 17 single-strain filtrates and the 23 community filtrates. The annotation gives the community-versus-single-strain contrast from a linear mixed-effects model fitting medium class as a fixed effect with focal-strain identity, plate nested within focal strain, and spent-medium identity as random effects: community filtrates reduced growth by 0.203 d_AUC_ units more than single-strain filtrates (df = 38.5, P = 2.2 × 10□□). B) Per-medium estimates (± SE) from a linear mixed-effects model fitting spent-medium identity as a fixed effect with focal-strain identity and plate as random effects. Each of the 40 media is contrasted against the unweighted mean of all 17 single-strain filtrates, which includes the medium itself when that medium is a single-strain filtrate; contrasts are corrected across media by Benjamini–Hochberg. Media are ordered by estimate and colored by class, and negative values indicate suppression exceeding the single-strain average. 35 of 40 media differed significantly from that average (q < 0.05), including 22 of 23 community filtrates. C) Mean d_AUC_ against spent-medium pH for the 39 filtrates in which pH was measured, spanning pH 5.8 to 8.0. The correlation is computed for the single-strain filtrates only (Spearman ρ = 0.35, P = 0.17, n = 17); community filtrates are plotted for context but were not tested, as their pH values did not differ detectably from one another (P = 0.48, ANOVA). Each spent medium was generated from a single conditioning culture, so replication within this experiment is technical: each focal strain was assayed on two 96-well plates, with the 17 single-strain filtrates present on both (six wells per focal strain by medium combination) and each of the 23 community filtrates present on one plate only (three wells).

Against the average of the 17 single-strain filtrates, 35 of 40 media differed significantly (q < 0.05; Figure 2B), including 22 of 23 community filtrates. The sole exception was Full - *B. thuringiensis* SC35 (estimate =-0.035, q = 0.31): removing SC35 alone erased most of the community suppression signal. The intact communities were the two most suppressive community filtrates (Full,-0.373; Full + *P. syringae* DC3000,-0.457), each exceeding every drop-out. The single most suppressive medium, however, was the *E. sibiricum* SC13 single-strain filtrate (-0.476), consistent with the null-model result below that community effects do not exceed their most inhibitory member (Figure 3). The next strongest single-strain filtrates were *B. thuringiensis* SC35 (-0.303) and *E. tasmaniensis* SC04 (-0.156); all others were above-0.20, weaker than 20 of the 23 community filtrates. Of SC35 and SC04, only SC35 appeared essential in community context: communities lacking SC04 remained strongly suppressive (Full - Gram-negative,-0.239), whereas both communities lacking SC35 were among the three weakest (Full - SC35,-0.035; Full - Gram-positive,-0.121). The two strongest single-strain suppressors, SC13 and SC35, shared no obvious growth, metabolic or biofilm phenotype (Supplementary Results).

**Fig 3.**
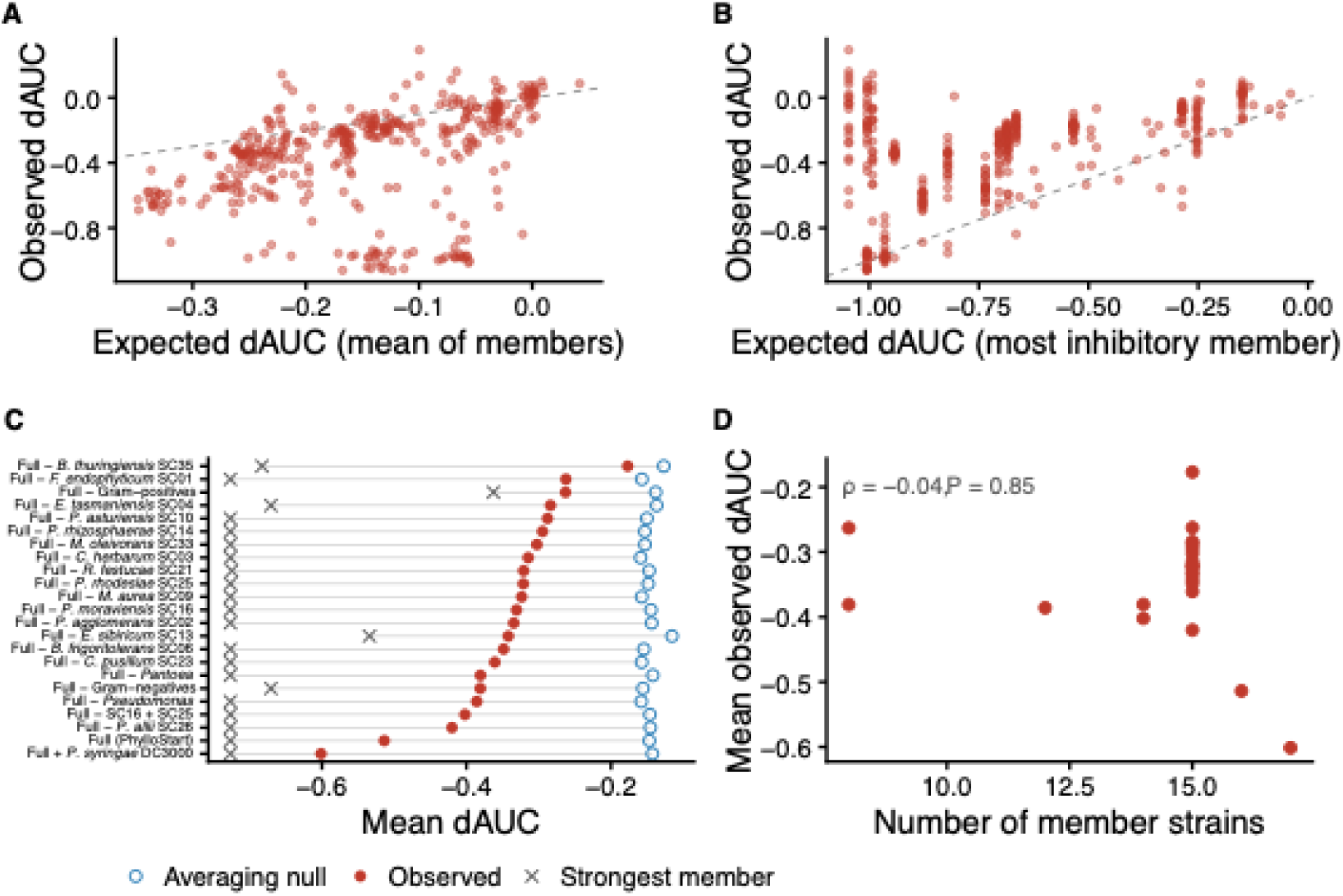
Community filtrate effects exceed additive expectations but not their most inhibitory member. For each of 23 community spent media and 17 focal strains (n = 391 combinations), the observed relative growth change (d_AUC_) is compared with two null expectations built from the single-species filtrates of that community’s own members. A) Observed d_AUC_ against the mean of member d_AUC_ values (additive/averaging null). Points below the diagonal indicate suppression stronger than expected from members in isolation. B) Observed d_AUC_ against the most negative member d_AUC_ (most-inhibitory-member null). Points above the diagonal indicate suppression weaker than the strongest member alone. C) Mean observed d_AUC_ per community (filled circles) with the corresponding averaging null (open circles) and most-inhibitory-member null (crosses); communities are ordered by observed effect. D) Mean observed d_AUC_ per community against the number of member strains. Values derive from the primary spent-media experiment, in which each medium was generated from a single conditioning culture and assayed in three wells (community filtrates) or six wells (single-strain filtrates) per focal strain; replication at this stage is technical.

Because pH modification can drive bacterial interactions [24], we measured filtrate pH against fresh medium and buffer (pH 6.8). Species identity had a significant effect on the pH of the spent media (P < 2.2 × 10^-16^, ANOVA). Single-strain filtrates ranged from pH 5.8 (*E. sibiricum* SC13) to 8.0 (*P. asturiensis* SC10; Figure 2C), whereas all drop-out community filtrates fell between pH 6.17 and 6.67 and did not differ from one another (P = 0.48, ANOVA): the pH extremes produced by individual strains were buffered in community context. Among single-strain filtrates, pH was not significantly correlated with mean d_AUC_ (Spearman ρ = 0.35, P = 0.17, n = 17; Figure 2C), and excluding *E. sibiricum* SC13, both the strongest acidifier and the strongest single-strain suppressor, weakened the association further (ρ = 0.22, P = 0.41). Conversely, *B. thuringiensis* SC35 changed pH only modestly (mean pH 6.5) yet was nearly as suppressive as SC13. Acidification therefore does not explain the suppressive effects of individual strains.

### Community effects exceed additive expectations but are weaker than their most inhibitory member

We compared each community filtrate with two null expectations built from its own members’ single-strain filtrates against the same focal strain (n = 391 community × focal strain combinations). Under the averaging null, a community’s d_AUC_ equals the mean of its members’ values; community filtrates were more suppressive than this in 321 of 391 combinations (median difference-0.10; Wilcoxon signed-rank P = 8.6 × 10□□³; Figure 3A), and the result was insensitive to weighting each member by its own growth in fresh medium (P = 2.9 × 10□□²; Supplementary Figure 11). Under the most-inhibitory-member null, a community equals its strongest member; community filtrates fell short of this in 349 of 391 combinations (median difference +0.31; P = 3.0 × 10□□□; Figure 3B). Every one of the 23 communities had a lower mean d_AUC_ than its member average, significantly so for 20 (P < 0.05; Figure 3C). Community effects are therefore emergent but bounded: stronger than the average of their members acting alone, but never stronger than their most inhibitory member.

These effects did not reflect the number of strains in the inoculum: neither observed suppression (Spearman ρ =-0.04, P = 0.85) nor departure from the additive null (ρ =-0.06, P = 0.78) was related to the number of member strains (Figure 3D), and the sixteen single-strain drop-outs, identical in size, spanned more than half the range of the entire panel. Which strains are present matters more than how many.

### Strain traits and relatedness are associated with interaction outcomes but do not predict them

We summarized each strain’s position in the 17 × 17 single-strain interaction matrix by its conditioning effect, the mean change in growth its filtrate imposed on the other strains, and its sensitivity, the mean change imposed on it by the other filtrates. The two were strongly negatively related (ρ =-0.75, P = 0.0007; Figure 4A): strains whose filtrates most reduced the growth of others were least affected by others’ filtrates, as expected if resource depletion rather than active antagonism dominates. Because the axes covary, we treat them as two readouts of a single strain-level property rather than as independent tests.

**Fig 4.**
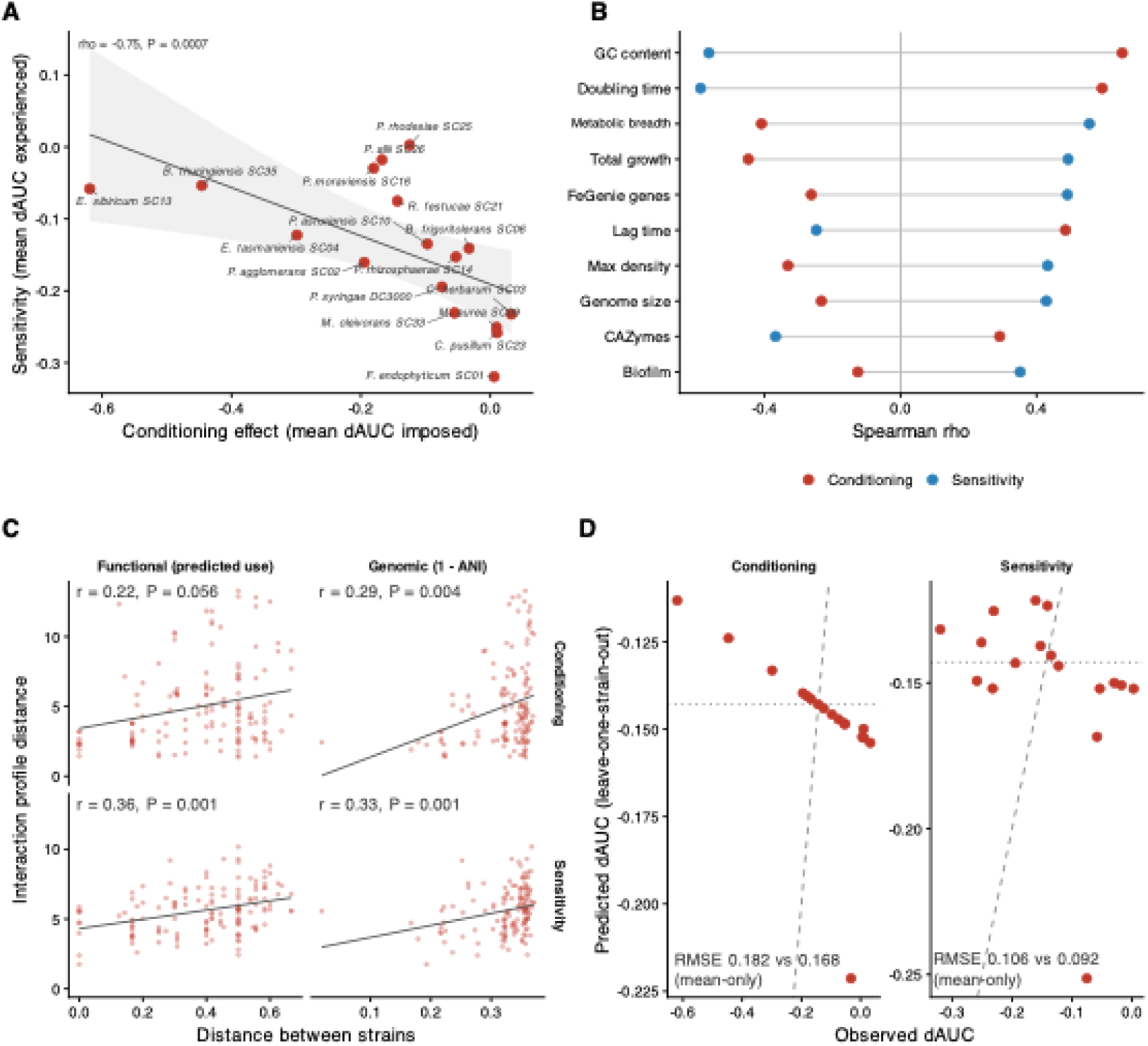
Strain traits and relatedness are associated with interaction outcomes but do not predict them. Each strain is summarized by two values from the 17 × 17 single-species spent-media matrix: conditioning effect, the mean d_AUC_ its filtrate imposed on all focal strains, and sensitivity, the mean d_AUC_ imposed on it by all filtrates. **(A)** The two axes against each other, one point per strain, with a linear fit and 95% confidence band. **(B)** Spearman correlation of ten genomic and phenotypic traits with each axis, shown as effect sizes. The panel shows the direction and relative magnitude of the associations across the strain panel; whether any trait explains variance in the assay beyond strain identity is tested separately at the level of individual wells (**Supplementary Table 5**). Metabolic breadth is the number of the 13 genomically scored EcoPlate substrates each strain is predicted to use. **(C)** Genomic distance (1 - ANI) and functional distance (Jaccard distance between genome-predicted substrate-use profiles) against interaction-profile distance, computed separately for conditioning and sensitivity because the interaction matrix is asymmetric; Mantel statistics from 9,999 permutations. **(D)** Observed against leave-one-strain-out predicted values from an elastic-net model on all ten traits, fitted on all seventeen strains. The dashed line is the 1:1 expectation and the dotted line the mean-only baseline; a root-mean-square error above the mean-only value indicates the model performs worse than predicting the mean. Conditioning and sensitivity values derive from the primary spent-media experiment, in which each medium was generated from a single conditioning culture and assayed in three wells (community filtrates) or six wells (single-strain filtrates) per focal strain; replication at this stage is technical.

Among ten genomic and phenotypic traits (Figure 4B), the strongest associations were with GC content, which is strongly conserved along broad phylogenetic lines, and growth strategy. Strains with higher GC content imposed less depletion on others (ρ = +0.65, P = 0.005) and were more affected by others’ filtrates (ρ =-0.56, P = 0.018), and longer doubling times behaved the same way on both axes (ρ = +0.59, P = 0.012 and ρ =-0.59, P = 0.013). Strains with greater genome-predicted metabolic breadth were less affected by others’ filtrates (ρ = +0.56, P = 0.021) and, more weakly, more depleting (ρ =-0.41, P = 0.103). Total growth and iron-acquisition gene content were related to sensitivity alone (ρ = +0.49, P = 0.044 and ρ = +0.49, P = 0.045), so greater iron-acquisition capacity did not translate into stronger conditioning of a shared medium (Supplementary Results). CAZyme repertoire, genome size and biofilm formation were unrelated to either axis (all P ≥ 0.086). Because the traits and the two axes are not independent, we report these associations as effect sizes without multiple-comparison correction and claim no individual association as significant. Metabolic breadth is also collinear with Gram status across this panel (ρ = +0.83), so a 17-member panel cannot separate metabolic capacity from broad taxonomy (Supplementary Results).

Relatedness showed similar structure (Figure 4C): genomic distance (1 - ANI) correlated with the distance between interaction profiles for both conditioning (Mantel r = 0.29, P = 0.004) and sensitivity (r = 0.33, P = 0.001). Functional distance, the Jaccard distance between genome-predicted substrate-use profiles, was marginally the stronger predictor of sensitivity (r = 0.36, P = 0.001) but was not significant for conditioning (r = 0.22, P = 0.056); the two directions were analysed separately because the interaction matrix is asymmetric (Supplementary Results). Relatedness therefore accounts for roughly a tenth of the variation in how strains interact.

None of these features supported prediction. An elastic-net model trained on all ten traits with leave-one-strain-out cross-validation performed worse than a mean-only baseline for both conditioning (root-mean-square error 0.182 against 0.168) and sensitivity (0.106 against 0.092), with negative cross-validated R² in both cases (-0.17 and-0.33; Figure 4D). Across 20 candidate mixed models (Methods), conditional R² was 0.432–0.436 for every model including the intercept-only null (Supplementary Table 5; Supplementary Figure 12); several trait models improved on the null by AICc, but total explained variance was unchanged, so traits redistributed variance already attributable to conditioner identity. In the null model, conditioner identity accounted for 32.7% of total variance and recipient identity for 10.0%, against 0.7% for plate. Strain traits and relatedness are thus associated with interaction outcomes but do not predict them; the associations that exist point to metabolic capacity and growth strategy rather than to annotated gene repertoires.

### Suppression of *P. syringae* requires the intact community

If the community protected plants by creating a chemical environment specifically hostile to *P. syringae*, the pathogen should be more sensitive to community filtrates than the commensals are. It was not. Averaged across the 23 community filtrates, *P. syringae* had a mean d_AUC_ of-0.319, ranking ninth of 17 focal strains in sensitivity and close to the commensal median (-0.324; range-0.912 for *B. frigoritolerans* SC06 to-0.065 for *P. rhodesiae* SC25). Its own filtrate was similarly ordinary, ranking tenth of 17 in conditioning effect (mean d_AUC_ = - 0.075). The pathogen thus sits within the commensal distribution on both axes, so the protection this community provides on plants is unlikely to be explained by the residual chemistry it leaves behind.

Because the *E. sibiricum* SC13 single-strain filtrate suppressed *P. syringae* almost completely, community suppression might simply reflect this one strain. It did not. *P. syringae* grew to a mean AUC of 0.51 in the full PhylloStart filtrate compared with 17.83 in fresh medium (n = 7 each, Mann–Whitney P = 3 × 10□□), but every one of the 21 drop-out filtrates permitted significantly more pathogen growth than the intact community (all Benjamini–Hochberg adjusted P = 0.008), including the 19 that retained SC13 (mean AUC 8.3 to 20.4; Supplementary Figure 13). The SC13 filtrate alone suppressed *P. syringae* as completely as the full community did (mean AUC 0.14 versus 0.51, P = 0.45), so the intact community matched but did not exceed its most inhibitory member. Suppression of the pathogen therefore depended on the full set of members and was lost by removing any one of them.

The full PhylloStart filtrate suppressed *P. syringae* far beyond its member average (d_AUC_ =-0.97 versus a null expectation of-0.21), but across the drop-out panel, pathogen suppression was not systematically greater than the additive expectation (P = 0.48), and no community exceeded the most inhibitory single member (*E. sibiricum* SC13, d_AUC_ =-0.99; P = 2.4 × 10□□). Suppression beyond the member average was thus specific to the intact community rather than a general feature of community subsets. pH alone also did not account for pathogen suppression. Across the 39 filtrates with pH measurements (pH 5.8 to 8.0), pH was not a significant predictor of *P. syringae* growth (Spearman ρ = 0.28, P = 0.09); the pathogen grew to within 10% of its fresh-medium growth in 15 filtrates spanning pH 6.2 to 7.9, filtrates indistinguishable in pH differed widely in suppression (Supplementary Results), and the intact community filtrates fall within a pH range that *P. syringae* tolerates. Relative growth changes were also reproducible in independently generated biological replicates of 22 community filtrates: across four focal strains including *P. syringae*, values correlated with those of the primary experiment (Spearman ρ = 0.21, P = 6.3 × 10□□), although the magnitude of suppression varied between batches (Supplementary Figures 14–18; Supplementary Results).

### Community context, not strain identity, shapes disease outcomes on tomato seedlings

Because spent-media assays remove the host, spatial structure and the nutrient landscape of the leaf, we asked whether in vitro outcomes could predict disease on plants. Seedlings were pre-inoculated with a single strain, a defined community (Supplementary Table 6) or buffer, and challenged three days later with *P. syringae* pv. *tomato* PT23 or buffer (Methods). We first defined when the assay resolves pathogen-induced disease: *Pst* challenged and unchallenged seedlings diverged from day 4–5, differed most at day 6 and converged after day 10, as seed reserves on water agar were exhausted (Figure 5A; Methods). Both experiments with a full time course showed this pattern (peak day 6; Experiment 1, P = 0.005; Experiment 2, P = 0.030), so all analyses use the mean score over days 5–9 (2–6 days post-challenge). Within this window, a single *P. syringae* challenge raised baseline-adjusted scores relative to buffer-only seedlings from +0.80 to +1.60 in Experiment 1 (P = 0.0064) and from +0.48 to +0.90 in Experiment 2 (P = 0.030).

**Fig 5.**
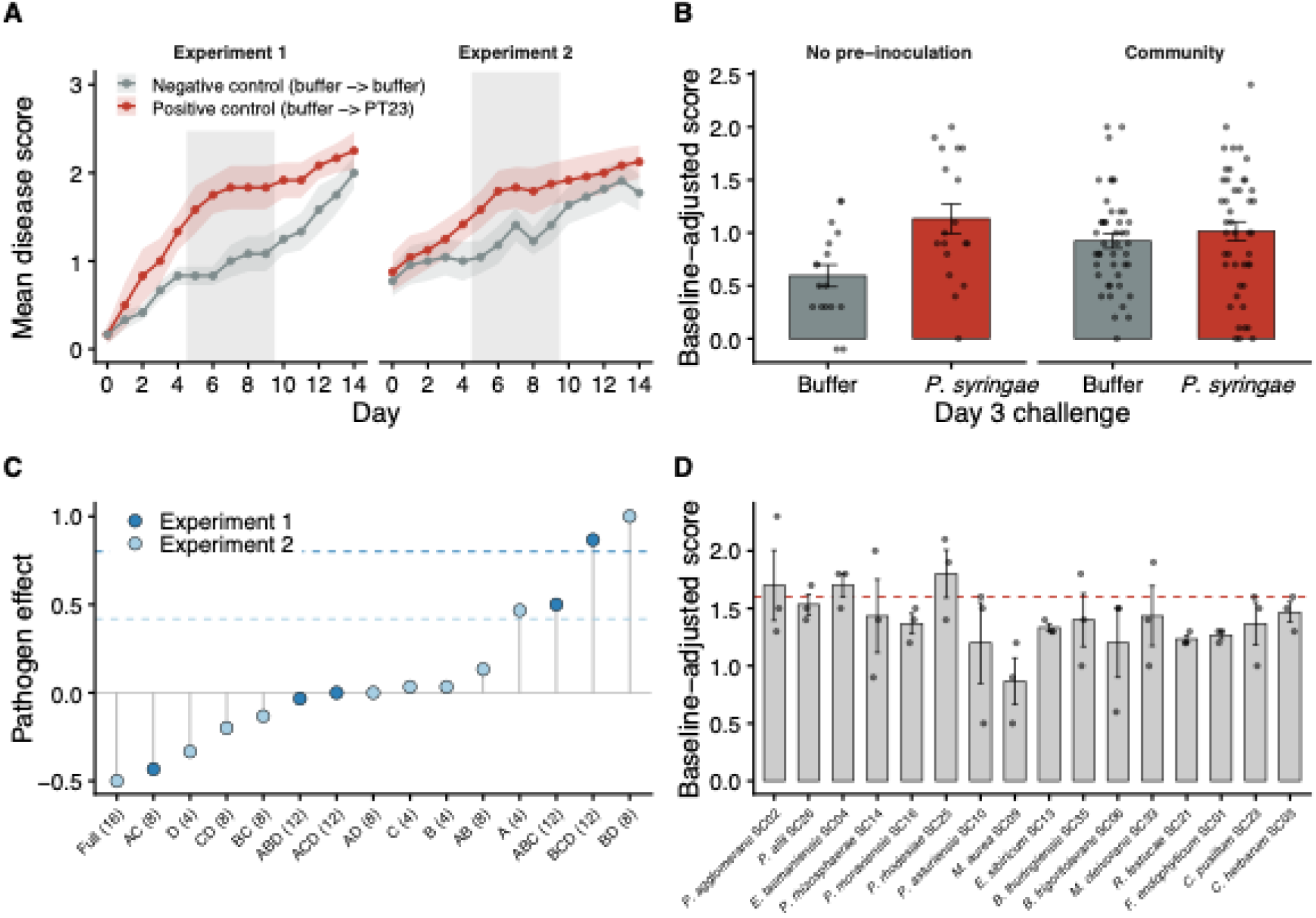
Community context, rather than the identity of individual strains, determines disease outcome on tomato seedlings. Tomato seedlings were inoculated on day 0 with a single commensal strain, a community, or buffer, challenged on day 3 with *P. syringae* pv. tomato PT23 or with buffer and scored daily for 14 days. Except in (A), the response variable is the baseline-adjusted disease score: the mean score across days 5–9 minus that seedling’s own day-0 score. (A) Daily mean disease score (± s.e.m.) for the two control arms of each experiment, negative control (no bacteria baseline) and positive control (pathogen reference). The shaded band marks the days 5–9 analysis window, bounded at its upper end by nutrient exhaustion in 1% water agar, after which scores in all arms converge. (B) Baseline-adjusted score by day-3 challenge, for seedlings receiving no pre-inoculation (left) and seedlings pre-inoculated with a community (right). Bars are means ± s.e.m. and points are individual seedlings (n = 3 per treatment). (C) Pathogen effect for each community, calculated as the mean baseline-adjusted score under *P. syringae* challenge minus the mean under buffer challenge. Fill color denotes the experiment; dashed lines mark the corresponding effect in that experiment’s no-pre-inoculation controls. Communities are combinations of four four-strain blocks: A, *Pseudomonas* (*P. asturiensis* SC10, *P. rhizosphaerae* SC14, *P. moraviensis* SC16*, P. rhodesiae* SC25); B, other Gram-negatives (*P. agglomerans* SC02, *E. tasmaniensis* SC04, *M. aurea* SC09, *P. allii* SC26); C, related Actinomycetota (*F. endophyticum* SC01, *C. herbarum* SC03, *R. festucae* SC21, *C. pusillum* SC23); and D, dispersed Gram-positives (*B. frigoritolerans* SC06, *E. sibiricum* SC13, *M. oleivorans* SC33, *B. thuringiensis* SC35). “Full” denotes PhylloStart, all 16 commensals, and the number in parentheses gives community richness. The four 12-member communities are single-block drop-outs: BCD is the community with the *Pseudomonas* removed and is compositionally identical to the Full-*Pseudomonas* spent medium in Figs 2 and 3, while AB and CD are the source communities for the Full-Gram-positive and Full-Gram-negative filtrates, respectively. (D) Baseline-adjusted score for seedlings pre-inoculated with each of the 16 commensal strains individually and challenged with *P. syringae*. Strains are ordered by their position in the core-gene phylogeny (Fig 1A); the dashed red line marks the mean of positive control. Replication is biological throughout. Experiment 1 tested all 16 commensals individually and five communities, with n = 3 seedlings per treatment and n = 6 per control arm; Experiment 2 tested the remaining ten communities in two batches, with n = 3 per treatment and n = 5-12 per control arm; Experiment 3 compared four carrier solutions with n = 6 per treatment.

Pre-inoculation with a community reduced this effect. Across all 15 communities (combinations of four four-strain blocks, A–D), tested over two experiments, the pathogen induced significant symptoms on bare seedlings but not on seedlings carrying a community (bare +0.54, community present +0.09; permutation P = 0.033; Figure 5B). Notably, pathogen-challenged bare seedlings and unchallenged community seedlings reached similar symptom levels (1.13 versus 0.92, P = 0.085), reflecting two distinct routes to mild symptoms: pathogen damage in the absence of a community, and a community-associated cost in the absence of the pathogen. Twelve of the 15 communities showed a smaller pathogen effect than the controls of the same experiment (Figure 5C), independent of richness (Spearman ρ =-0.08, P = 0.79); the six eight-member communities alone spanned-0.43 to +1.00. Pre-colonization was not uniformly protective: BD (+1.00 against +0.42) and BCD (+0.87 against +0.80) clearly, and A (+0.47 against +0.42) marginally, showed a pathogen effect at or above their experiment’s positive (pathogen only) control level. In contrast, no single strain altered disease: scores did not differ among the 16 commensals tested singly (Kruskal–Wallis P = 0.72), and none differed from the pathogen-alone control (all P > 0.19; Figure 5D), so strain-level suppressive phenotypes in spent media had no counterpart in seedling disease. A community-level comparison across the two assays is limited by design: the spent-media panel (the full community and its single-strain and single-block drop-outs) and the seedling panel (four-strain blocks and their combinations) share only two compositionally identical communities, the full PhylloStart community and the Pseudomonas-free community (seedling BCD, equivalent to the Full-Pseudomonas filtrate). Both assays nonetheless converge on the same result: pathogen suppression required the intact community and was not attributable to any single member.

Pre-inoculation with a community was not entirely benign: pooled across communities, seedlings receiving a community without the pathogen scored modestly higher than untreated seedlings (+0.92 versus +0.59 baseline-adjusted, P = 0.010; Figure 5B), an effect independent of the inoculation carrier that did not survive correction at the level of individual communities (Supplementary Figure 19; Supplementary Results). Pre-colonized seedlings nonetheless showed less disease under pathogen challenge than sterile seedlings, so this community-associated cost does not account for the suppression result. Disease outcome after pathogen challenge therefore depends on the composition of the resident community rather than on any single member, and community pre-colonization itself carries a cost to the seedling that varies with composition.

## Discussion

Harnessing microbiomes for host or environmental health requires understanding how community composition shapes function [25]. Interspecific interactions govern the assembly, stability and function of microbiomes [1, 26], yet they remain difficult to predict from genomic features or from members studied in isolation, which limits our ability to leverage plant microbiomes for plant health [27]. Using a 16-member tomato phyllosphere SynCom [20] and the pathogen *P. syringae*, we tested whether genomic and phenotypic features predict pairwise and/or community-level interaction outcomes, and how these two types of interactions might differ. We found that community effects were emergent but bounded: community filtrates were consistently more suppressive than the average of their members acting alone, yet never exceeded their single most inhibitory member (**Figure 3**). Suppression of *P. syringae* followed the same logic: the intact community suppressed the pathogen far beyond its member average, but every drop-out filtrate permitted more pathogen growth, including those that retained the strongest single suppressor, *E. sibiricum* SC13 (**Supplementary Figure 13**).

On seedlings, protection likewise required a community and was unrelated to community richness (**Figure 5**). The seedling data cannot test the bounded half of this result, because no single commensal altered disease and there is no most-inhibitory member to compare against. They do, however, reproduce the other half: suppression emerged only at the community level. This convergence is consistent with colonization resistance being a community-level property rather than the sum of its members: in defined human gut communities, individual symbionts provided negligible protection, whereas assembled communities restricted pathogens by orders of magnitude, depending on nutrient-use overlap between community and pathogen [16]. Our drop-out panel recovers that structure in the phyllosphere, with two key differences. First, *P. syringae* was indistinguishable from the commensals in the cell-free assay, so the resistance we measured in vitro is a general constraint on invasion into an already-conditioned niche rather than a pathogen-specific one [14]. Second, whereas assembled gut communities exceeded single member effects on the pathogen, our communities matched but did not surpass the effect of their strongest member. This may be expected if the dominant currency is resource pre-emption rather than synergistic antagonism, and is consistent with the absence of strong higher-order positive effects among culturable bacteria [28].

Overall, we find that suppression observed in single-strain assays does not generally translate to community contexts, suggesting the performance of candidate biocontrol agents is likely to depend strongly on the other members present. While *E. sibiricum* SC13 acidified its filtrate below pH 6 and was the most suppressive single strain, the community lacking it remained strongly suppressive across focal strains. In contrast, removing *B. thuringiensis* SC35, which reduced growth without modifying pH, erased most of the community signal (**Figure 2B**). Community filtrates converged on a narrow pH range relative to single strains, and pH explained neither single-strain suppression nor suppression of the pathogen (**Figure 2C**). Communities therefore buffer the environmental extremes their members generate in isolation, suggesting that environmental modification may be a particularly poor trait to select for when engineering microbiome function. SC35 carries 13 biosynthetic gene clusters, including clusters for zwittermicin A and the siderophores bacillibactin and petrobactin [29–31], making iron competition a plausible contributor to suppression, as siderophore-mediated iron competition predicted both pathogen suppression and protection of tomato plants across 2,150 rhizosphere isolates [32], and our assay retains secreted siderophores in an iron-limited medium, although we cannot distinguish it from other diffusible mechanisms here.

The trade-off between conditioning and sensitivity (**Figure 4A**), together with associations with growth strategy and genome-predicted metabolic breadth (**Figure 4B**), points to competition for shared resources rather than active antagonism as the dominant interaction. Relatedness showed weak but detectable structure (**Figure 4C**), consistent with phylogenetic clustering of interactions among isolates from polymicrobial infections [33] and synthetic gut consortia [18]. Yet no trait set outperformed a mean-only baseline, and traits redistributed variance already attributable to strain identity (**Figure 4D**), echoing the weak transfer of culture-derived genomic predictors to performance in complex environments [7]. Community assembly has been predicted from genome-inferred resource use in soil bacteria [34], but in single-carbon-source minimal medium, which collapses the resource landscape onto one axis. In a complex medium such as King’s B, partitioning, cross-feeding and environmental modification operate at once, and future assays performed in defined single-substrate media would be necessary to test whether predictability depends on resource-landscape breadth. Likewise, curated genome-scale metabolic models of *Arabidopsis* leaf isolates recapitulated interaction outcomes in planta [35], but they reconstruct strain-specific resource use mechanistically, whereas our traits are additive summaries. More importantly, their unit of prediction is the pair, and our central result is that community-level effects are not recoverable from member-level information, which mirrors the same limitation our own additive and most-inhibitory-member nulls encountered. On plants, no single commensal altered disease despite their wide range of suppressive phenotypes in vitro. Our strains were chosen for taxonomic breadth rather than screened for biocontrol activity, and in a screen of 224 Arabidopsis leaf isolates only about 10% conferred full protection against *P. syringae* pv. *tomato* DC3000 [36].

Our interpretations are subject to the limits of in vitro spent-media assays. Filtrates exclude contact-dependent mechanisms and were collected at stationary phase to standardize resource exhaustion across strains with very different growth rates, a point at which the metabolic regime is poorly characterized and need not resemble what a colonizer meets on a leaf. King’s B and the absence of spatial structure further limit realism, and we did not manipulate iron availability, redox state or cofactor limitation, so the interactions we report are specific to these conditions. Our seedling experiments provide the ecologically realistic counterpart but measure disease rather than the chemistry underlying it. Pathogen load could not be quantified because community members cannot be resolved on plates when grown together. The magnitude of community suppression of *P. syringae* also varied between independently generated batches of filtrates, although the ranking of filtrates was preserved (**Supplementary Figures 17-18**), and only about half of all genes could be functionally annotated, constraining every annotation-derived predictor. Finally, our community was assembled to represent the family-level diversity of the tomato phyllosphere and includes several closely related members; a panel built to maximize phylogenetic spread would likely show a weaker relatedness signal and a different distribution of interaction strengths. Defining a phyllosphere-mimicking medium and repeating these assays across controlled nutrient and iron regimes is, in our view, the most important next step, alongside nutrient supplementation, fractionation, metabolomics and spatially structured co-culture to identify the metabolites mediating suppression.

Together, our results show that genomic and phenotypic traits define moderately conserved strategies across lineages but are poor predictors of interaction strength, and that in vitro no community lacking any one of its members reproduced the pathogen suppression of the full PhylloStart community. In planta, disease suppression was similarly reliant on community presence and single strain pre-inoculation was not sufficient to suppress pathogenicity. Community-level suppression instead arose from collective activity, likely reflecting complementary resource use and the accumulation of multiple diffusible metabolites. Engineering phyllosphere communities for plant protection will therefore benefit from manipulation of whole-community composition rather than targeting single taxa, especially when effects are measured in isolation.

## Materials and methods

### Strain Identifications

We used the 16 members of the tomato-derived SynCom PhylloStart, eight Gram-negative and eight Gram-positive strains whose isolation and composition are described in [20], and the model plant pathogen *Pseudomonas syringae* pv. *tomato* DC3000 and PT23 [20]. PT23 was used only for seedling challenge and was not sequenced or included in the spent media assays; genomic and in vitro analyses therefore cover 17 isolates. Commensal members were isolated from tomato leaves and share a co-occurrence history on this host [20]; closely related members were retained deliberately so that the influence of relatedness on interactions could be tested.

Culture identity and purity were confirmed by Sanger sequencing of the near full-length 16S rRNA gene (primers 27F/1492R; **Supplementary Methods**) and BLAST against the 16S rRNA database [37]. BTyper3 was used to resolve the species identity of *Bacillus* SC35 [38, 39].

### Whole genome sequencing of bacterial isolates

Genomic DNA was extracted from 2–3 day KB cultures. Short-read libraries (Qiagen DNeasy Blood and Tissue Kit) were sequenced at the California Institute for Quantitative Biosciences (UC Berkeley), and long-read extraction (MagMAX Microbiome Ultra Kit) and Oxford Nanopore sequencing (SQK-NBD114 native barcoding) were performed by SeqCoast Genomics (Portsmouth, NH, USA; **Supplementary Methods**).

### Hybrid genome assembly

Short read quality was assessed with FastQC [40], MultiQC [41] and NanoPlot [42]. Short reads were trimmed with FastP [43], long reads were filtered with Filtlong, and hybrid assemblies were generated with Unicycler [44]. Putative plasmids were assembled and identified using plasmidSPAdes [45]. Genome assembly statistics were summarized with Quast (**Supplementary Table 2**) [46] and genome completeness and contamination were assessed with CheckM (**Supplementary Table 3**) [47]. Average nucleotide identity between genomes was calculated with OrthoANI [48].

### Annotation of genomic data

Genomes were annotated with Prokka [49], KEGG pathways with BlastKOALA [50], and COG categories with EggNOG [51]. Orthologous genes were identified with OrthoFinder for the 16 commensal genomes and for all 17 genomes [52]. Secondary metabolite biosynthetic gene clusters were identified with antiSMASH [53], iron-related genes with FeGenie [54], carbohydrate-active enzymes (CAZymes) with dbCAN3 [55–57], and metabolic potential with gapseq [58] and gapmind [59].

### Phylogenetic analyses

Phylogenies were inferred from the 16S rRNA gene using ape [60], and from 107 conserved single-copy genes using bcgTree [61, 62] with hmmsearch [63], MUSCLE and Gblocks [64, 65], and RAxML with 100 bootstraps [66]. The core-gene phylogeny shown in Fig 1A includes all 17 strains; the comparison of 16S and core-gene topologies in Supplementary Figure 1 is restricted to the 16 PhylloStart members.

### Growth curve measurements in sterile media

KB cultures grown for 24–72 h at 28.0°C were normalized to an optical density at 600 nm (OD600) of 0.002, and 5 μl was inoculated into sterile KB in 96-well plates at 28.0°C. OD600 was read every 12 h for 72 h on a Tecan spectrophotometer after 5 s of orbital shaking.

### Assaying biofilm formation by staining

Biofilm formation was quantified by crystal violet staining of 100-fold diluted two-day KB cultures grown without shaking in flat-bottom 96-well plates at 28.0°C for 3 days, read at OD600 after solubilization in 95% ethanol (Supplementary Methods).

### Spent (conditioned) media generation

Single-strain spent media (n = 17) were generated by inoculating one colony from a KB agar plate into 40 ml of KB broth, and community spent media (n = 23; **Supplementary Table 4**) by inoculating one colony of each member into the same volume. Cultures were incubated for 48 h at 28.0°C, by which time all strains reach stationary phase (**Supplementary Figure 6**), then centrifuged (10 min, 4,000 RCF), passed through a 0.22 μm filter and stored at-20.0°C.

### Measuring pH of spent media

Filtrate pH was measured with Whatman indicator strips (Supplementary Methods) and compared among filtrates by one-way ANOVA, separately for single-strain and community filtrates because the latter are largely drop-outs of a common parent community and are not independent. Each single-strain filtrate’s mean pH was correlated with its mean d_AUC_ across the 17 focal strains (Spearman’s rank correlation); community filtrates were not tested because their pH did not differ detectably.

### Experimental growth curves in spent media

Focal strains were grown for 48 h in KB, washed twice in sterile MgCl□, normalized to OD600 = 0.002 (Tecan Infinite F50), and 5 μl was inoculated into wells containing each of the 40 spent media. Each focal strain was assayed on two 96-well plates, with six wells per single-strain filtrate and three per community filtrate, and growth was read as OD600 every 12 h for 72 h.

To test reproducibility, we generated three new biological replicates of 22 of the 23 community spent media (all except Full-SC16&SC25) and grew *P. syringae* DC3000, *P. rhizosphaerae* SC14, *P. moraviensis* SC16, and *R. festucae* SC21 in all 66, randomized within a 96-well plate with fresh-medium and negative controls and measured as above.

### Total growth in spent media calculations and comparisons

Growth curves were analyzed with gcplyr in R [67, 68]. Area under the curve (AUC) was used as total growth, and relative growth change (d_AUC_) was calculated by contrasting growth in spent (SM) and sterile (ST) medium, following Weiss et al., who termed it an inhibition factor [18]; negative values indicate reduced growth. We use growth change rather than inhibition because this design does not distinguish reduced growth arising from resource depletion during conditioning from that arising from active antagonism. d_AUC_ was visualized with pheatmap [69].

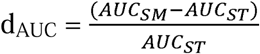

Because each focal strain contributed 192 wells across two plates, wells were not treated as independent. Linear mixed models were fitted in lme4 [70], with p-values from lmerTest [71] and contrasts from emmeans [72]. Community and single-strain filtrates were compared by fitting spent-media class as a fixed effect, with focal strain, plate nested within focal strain, and spent-media identity as random effects, and cross-checked with a paired test on strain-level means (n = 17). For per-medium estimates, spent-media identity was fitted as a fixed effect, each medium was contrasted against the mean of the single-strain filtrates, and p-values were adjusted by the Benjamini–Hochberg procedure. Because each community filtrate was assayed on one plate, a systematic plate effect was tested using fresh-medium controls run on both plates for every focal strain. Model diagnostics are given in Supplementary Figure 9, and the rationale for the random-effects structure in the Supplementary Methods.

To test whether any trait explained variance in d_AUC_ beyond strain identity, we fitted a pre-specified set of 20 maximum-likelihood mixed models in lme4 against an intercept-only null; the model set and its shared random-effects structure are given in the Supplementary Methods. Marginal and conditional R² were computed with r.squaredGLMM in MuMIn [73].

### Resource utilization with Biolog plates

We investigated resource utilization using Biolog Ecoplates (Biolog Inc., Hayward, CA, USA). Washed 3-day KB cultures (four pooled replicates) were normalized to OD600 = 0.02 and inoculated into EcoPlates, which were read at OD590 and OD750 over 3 days and again at day 7 to confirm that growth had plateaued (**Supplementary Methods**). Readings were corrected against the control well. We measured the average well color development (AWCD) to quantitatively evaluate the Biolog results using the spectrophotometer measurements [74, 75].

For replicate *j* at time *t,* the AWCD is given as:

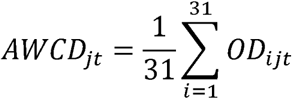

Here, OD*_ijt_* denotes the corrected OD for well *i* of replicate *j* measured at time *t.* Therefore, the adjusted OD can be measured with OD = OD*_ijt_*/AWCD*_jt_* and these adjusted values were used for all analyses and interpretations.

### Community sequencing with 16S rRNA gene amplification

Pellets from 2 ml of each community culture (**Supplementary Table 4**) were stored at - 70.0°C and sent to SeqCoast Genomics, where DNA was extracted (Qiagen DNeasy 96 PowerSoil Pro QIAcube HT Kit with 40 min bead beating), and the V3–V4 region of the 16S rRNA gene was amplified (Zymo Quick-16S Plus NGS Library Prep Kit; primers 341f/806r) and sequenced on an Illumina NextSeq2000 (2 × 300 bp). Full protocols are given in the Supplementary Methods. Remaining adapters and primers were trimmed with Cutadapt [76], and amplicon sequence variants (ASVs) were inferred with DADA2 using default parameters [77, 78], classified against SILVA [79], and analyzed with phyloseq [80].

### Tomato seed sterilization and germination

Seeds of organic tomato cv. Moneymaker were surface-sterilized with ethanol and sodium hypochlorite (Supplementary Methods) and germinated singly in 15 ml tubes of 1% water agar at 21.0°C in darkness for about 5 days, then moved to a growth chamber (24.0°C, 15 h light: 9 h dark). Seedlings were 12 days old at first inoculation and remained in the same tubes, without added nutrients, for the duration of the experiment.

### Seedling disease assays

Seedling inoculations followed published methods [81]. Each seedling received a first inoculant at day 0 and a challenge at day 3, with four control schemes: buffer then buffer (untreated), buffer then *P. syringae* (pathogen challenge; the reference for all pre-inoculation treatments), *P. syringae* then *P. syringae*, and *P. syringae* then buffer.

Inocula were prepared from 24–48 h KB cultures, washed and resuspended in 10 mM MgCl□, and normalized to OD600 = 0.002 (higher densities caused rapid necrosis in preliminary trials) in 10 mM MgCl□ with 0.015% (v/v) Silwet L-77; 10 µl was applied to the seed leaves, and controls received carrier alone (Supplementary Methods). *P. syringae* pv. *tomato* PT23 rather than DC3000 was used for challenge to maintain comparability with seedling and whole-plant protection assays established in this system [3].

Three experiments are reported. Experiment 1 tested all 16 commensals individually against *P. syringae* challenge, together with five communities against both challenges (n = 3 seedlings per treatment, n = 6 per control arm). Experiment 2 tested the remaining ten communities against both challenges in two batches (n = 3 per treatment, n = 5–12 per control arm). Experiment 3 compared four carrier solutions (n = 6 per treatment). Community membership by experiment and the carrier solutions are listed in Supplementary Table 6 and the Supplementary Methods.

### Disease scoring and analysis

Disease was scored daily from day 0 to day 14 following [81]: no symptoms (0), mild speckling (1), extensive speckling and/or chlorosis (2), and leaf necrosis and/or leaf detachment (3), with half-point increments. Tubes were oriented with labels facing away from the scorer, and seedling positions were randomized.

The analysis window was defined from the control arms alone, as the interval over which pathogen-challenged and untreated seedlings differed: days 5–9 of scoring (2–6 days post-challenge), applied unchanged to all treatments. Later timepoints were excluded because seedlings on 1% water agar exhaust their seed reserves and develop stress symptoms, such as chlorosis, independently of inoculation.

Each seedling’s day-0 score was subtracted from its day 5–9 mean, and baseline-adjusted values are reported throughout. One negative-control seedling in Experiment 2 scored 3 at day 0 and was excluded as non-viable; re-including it does not change any conclusion (interaction P = 0.021; community-alone effect P = 0.005). Group comparisons used Mann–Whitney U and Kruskal–Wallis tests, and the pathogen × pre-inoculation interaction was tested by permuting challenge assignment within pre-treatment class and experiment (20,000 permutations). Null expectations for community filtrates were computed per community from the single-strain d_AUC_ values of its own members against the same focal strain, as the member mean (averaging expectation) or the most negative member value (most-inhibitory-member expectation), and compared with observed values by Wilcoxon signed-rank tests across all community × focal strain combinations and within each community. Multiplicative (Bliss independence) expectations were not used because member contributions to shared resource depletion are not independent (Supplementary Methods).

### Statistical analyses, modeling, and visualization

All statistical analyses were performed in R using Rstudio [67]. The community sequencing data was statistically analyzed using the *vegan* package [82]. Data visualization was performed with *ggplot2* [83], *ggpubr* [84], *RColorBrewer* [85], and *forcats* [86]. Phylogenetic trees were visualized using a combination of the iTOL: Interactive Tree of Life v6 and *ggtree* in R [87, 88]. We used elastic-net regression with leave-one-strain-out cross-validation in R with *glmnet*. Model performance was compared to a null mean-only baseline.

## Data availability statement

Raw whole-genome sequencing data are available at the NCBI SRA under BioProject PRJNA1297335 (long-read) and PRJNA1374443 (short-read), and 16S rRNA gene amplicon sequencing of the spent-media communities under PRJNA1297786. BioSample Accessions for the hybrid assembled genomes are given in Supplementary Table 1. Data underlying the figures are deposited on Dryad (https://doi.org/10.5061/dryad.v6wwpzh9z): spent-media growth assays (raw and mean relative growth change for the primary and replicate experiments; Figs 2–4), spent-media pH measurements (Fig 2C), and tomato-seedling disease scores (Fig 5). Genome assembly statistics, synthetic-community compositions, and the mixed-model selection results are provided in Supplementary Tables 2–6.

## Supporting information

Supplementary Methods, Supplementary Results, Supplementary Figure

## Acknowledgements

This research used the Savio computational cluster resource provided by the Berkeley Research Computing program at the University of California, Berkeley (supported by the UC Berkeley Chancellor, Vice Chancellor for Research, and Chief Information Officer).

