## Supplementary Methods, Supplementary Results, Supplementary Figure for "Community context supersedes species identity in bacterial interactions and disease outcomes in the tomato phyllosphere"

This section gives protocol details summarized in the main-text Materials and Methods.

**Strain identification and culture purity**

To validate culture purity, we sequenced and amplified the 16S rRNA gene using the forward 27F 5’-AGAGTTTGATCCTGGCTCAG-3’ and reverse 1492R 5’-GTTACCTTGTTACGACTT-3’ primers. Polymerase chain reaction (PCR) was performed using bacterial culture as DNA template that was diluted 1:200 in MgCl2 buffer. Bacteria were grown from frozen stock in King’s B (KB) liquid medium for 48 hours at 28.0°C. We performed 20 μl reactions using Platinum HotStart Master Mix (Invitrogen) with the following cycle conditions: initial denaturation at 94.0°C for 3 min followed by 30 cycles of denaturation at 94.0°C for 30 sec, annealing at 48.4°C for 30 sec, and extension at 72.0°C for 1 min 30 sec with a final extension at 72.0°C for 10 sec.

The UC Berkeley DNA Sequencing Facility performed PCR clean up and Sanger sequencing on the amplified products (https://ucberkeleydnasequencing.com/home). The chromatograms were visualized and sequences extracted using 4peaks (https://nucleobytes.com/4peaks/) and were identified through BLAST analysis against the 16S rRNA ribosomal sequences database [S1]. For the *Bacillus* strain SC35, we used BTyper3 to precisely identify species identity within the *Bacillus* genus [S2, S3].

**Whole genome sequencing of bacterial isolates**

The bacterial genomes of all isolates were sequenced using long and short read technologies. Before DNA extraction, the bacteria were grown from frozen glycerol stocks in KB liquid media in a shaking incubator at 28.0°C. After 2-3 days of growth, the bacteria were spun down at 4,000 RCF for 10 minutes. After centrifugation, the supernatant was removed, and the pellet was used as input for DNA extraction. DNA extractions for short read sequencing were performed with Qiagen DNeasy Blood and Tissue Kits and short read sequencing was performed at the California Institute for Quantitative Biosciences at the University of California Berkeley (Berkeley, CA). Long-read DNA extraction and sequencing was performed by SeqCoast Genomics (Portsmouth, NH, USA) using Oxford Nanopore Technologies. DNA was extracted using the MagMAX Microbiome Ultra Nucleic Acid Isolation Kit which uses bead beating lysis. DNA samples were prepared for whole genome sequencing using the Oxford Nanopore Technologies SQK-NBD114 native barcoding kit with Long Fragment Buffer to promote longer read lengths.

Long reads were filtered with Filtlong (https://github.com/rrwick/Filtlong); long-read quality was assessed with NanoPlot from the NanoPack software suite [S4].

**Phylogenetic analyses**

We visualized the phylogenetic relationships of the bacteria in this study (16 SynCom members and the *P. syringae* pathogen) using different sets of genetic information, as different segments or sets of DNA may reveal non-congruent phylogenetic histories. First, we evaluated the relationships of the 16 SynCom members using the 16S rRNA gene sequence and visualized them using the Analyses of Phylogenetics and Evolution (ape) package with R [S5]. The comparison of 16S and core-gene topologies in Supplementary Figure 1 is therefore restricted to these 16 strains, whereas the core-gene phylogeny in Figure 1A includes all 17. Next, we used BCGtree to reconstruct the phylogenetic relationships of the 17 bacteria using a set of conserved prokaryotic genes previously described by Dupont et al. (2012) [S6, S7]. A proteome fasta file for each of the bacterial species was used as input for bcgTree to search for the 107 conserved single copy genes using hmmsearch [S8]. The conserved genes were aligned using muscle and the alignment was polished using gblocks [S9, S10]. The polished conserved gene alignment file was used as input for tree building with RAxML with 100 bootstrap searches and visualized as previously described [S11].

**Biofilm formation**

To assess biofilm capabilities of each of the strains, we performed biofilm staining with crystal violet. Bacteria were grown individually in KB liquid media from frozen stock and incubated at 28.0°C for 2 days. The bacteria were diluted 100-fold into sterile KB media in flat-bottom 96 well plates. The plates were incubated without shaking at 28.0°C. Following incubation, the planktonic cells and media were discarded, and the plate was rinsed with water twice. After rinsing, 0.1% crystal violet was added to each well and incubated at room temperature for 15 minutes. The crystal violet was rinsed out as previously described. The plate was allowed to dry overnight at room temperature before the crystal violet was solubilized with 95% ethanol. The amount of stain was recorded by taking OD600 measurements with the spectrophotometer.

**Spent (conditioned) media generation**

Individual bacteria were streaked onto KB medium hard agar plates from frozen stock with 10 μl inoculating loops. The plates were incubated for 24-72 hours at 28.0°C. Monoculture spent media was started by inoculating a single colony into 40 ml of KB liquid medium (n = 17 monoculture, or single species, spent media types). To generate community spent media, individual colonies of each respective bacteria were picked and inoculated into 40 ml of KB liquid medium (n = 23 community spent media types, Supplementary Table 4). All liquid cultures were incubated at 28.0°C for 48 hours which was sufficient time for the cultures to be visibly turbid. Because all species reach stationary phase by 48 hours (Supplementary Figure 6), we did not expect the difference in initial starting concentration (1 colony vs. 8-17 colonies for community spent media) to impact the comparisons. Following incubation, the culture was prepared for filtration by centrifugation for 10 min at 4,000 RCF in an Eppendorf centrifuge model 5425. After centrifugation, cells and debris were removed through a 0.22 μm filter and the flow-through was collected into sterile tubes and stored in -20.0°C freezers.

**Measuring pH of spent media**

The pH of the sterile and spent media was measured using Whatman pH indicator strips. Both broad and narrow range strips were used. After 48 hours of growth, the pH of the bacterial cultures was measured by dipping an indicator strip into the well for all replicates, first with the broad range strip. If the measured pH was within the range of the narrow strip, we also measured the well using the narrow range strip for greater precision.

**Growth curves in spent media**

To prepare focal bacteria, each isolate was revived from frozen stock in KB liquid medium and incubated for 48 hours at 28.0°C. After incubation, the cultures were centrifuged for 10 minutes at 4,000 RCF and resuspended in sterile MgCl2 buffer twice to remove their own residual media. Inocula were prepared by normalizing the resuspended bacteria to an optical density (OD600) of 0.002 measured by a Tecan spectrophotometer (Infinite F50). The 40 different spent media types were each aliquoted across 96-well plates for experimental testing. Growth curves were started by inoculating each spent media well with 5 μl of normalized bacteria.

**Resource utilization with Biolog EcoPlates**

Each bacteria was grown for 3 days in 4 replicate cultures of KB liquid media at 28.0°C with 150 RPM. After 3 days, the supernatant was removed by centrifugation, and the bacteria were resuspended with 1 ml of MgCl and concentrated into a single tube. The bacterial density was normalized to a starting OD600 of 0.02 before inoculation into the Ecoplate. The Ecoplates were read at OD590 and OD750 three times the first day, two times the second and third day, and then again at seven days to establish that they had indeed reached carrying capacity by the end of the third day (which was found to be true for each species tested). To interpret the Biolog results we first corrected the OD measurements by subtracting the OD values of the control wells from the wells containing substrates.

**Community 16S rRNA gene amplicon sequencing**

To prepare community samples for 16s rRNA gene sequencing, we centrifuged 2 ml of each community (Supplementary Table 4) to obtain a pellet. Bacterial pellets were stored in the -70.0°C freezer prior to being shipped to SeqCoast Genomics (Portsmouth, NH, USA). Samples were transferred to MagMAX Microbiome Bead Beating Tubes (FisherSci #A42351) and then mixed with Qiagen's CD1 Lysis Buffer (from the Qiagen DNeasy 96 PowerSoil Pro QIAcube HT Kit [#47021]). Bead beating was then performed using a Vortex Genie 2 for 40 minutes. For samples intended for short-read or amplicon sequencing, our standard bead-beating time is 40 minutes. After bead-beating lysis, samples were extracted using the Qiagen DNeasy 96 PowerSoil Pro QIAcube HT Kit (#47021).

DNA samples were prepared for 16S V3/V4 amplicon sequencing using the Zymo Quick-16S Plus NGS Library Prep Kit (#D6421, https://www.zymoresearch.com/products/quick-16s-plus-ngs-library-prep-kit-v3-v4-udi), which uses unique dual indexes. To amplify the 16s rRNA gene, the 341f (CCTACGGGDGGCWGCAG, CCTAYGGGGYGCWGCAG, 17 bp) and 806r (GACTACNVGGGTMTCTAATCC, 24 bp) primers were used. The forward primer 341f is a mixture of the two sequences listed. The amplification PCR conditions were as follows: initial denaturation at 95.0°C for 10 min followed by 35 cycles of denaturation at 95.0°C for 30 sec, annealing at 55.0°C for 30 sec, and extension at 72.0°C for 3 min with a final extension at 72.0°C for 6 min.

Sequencing was performed on the Illumina NextSeq2000 platform using a 600-cycle XLEAP-SBS flow cell kit to produce 2x300bp paired reads. A 30-40% PhiX control (unindexed) was spiked into the library pool to support optimal base calling of low diversity libraries on patterned flow cells. Read demultiplexing, adapter trimming, and run analytics were performed using DRAGEN v4.2.7, an on-board analysis software on the NextSeq2000.

To analyze the community sequencing reads, we used Cutadapt to trim any remaining adapter and primer sequences from the demultiplexed reads [S12]. We performed marker gene analysis in R using the DADA2 package using default parameters for filtering and trimming [S13, S14]. DADA2 was also used for dereplication, merging paired reads, and chimera removal. The resulting ASVs from DADA2 were matched to the SILVA database. The output along with sample metadata were combined into a phyloseq object for downstream statistical analyses [S15].

**Tomato seed sterilization and germination**

Seeds of organic tomato cv. Moneymaker were surface-sterilised in batches. Seeds were agitated in approximately 30 ml of 70% ethanol for 1 min, the ethanol decanted, and the seeds transferred to 15 ml of 2.75% sodium hypochlorite supplemented with 20 µl of 0.2% (v/v) Tween 20 and shaken for 20 min. Seeds were then rinsed four times in 45 ml of sterile MilliQ water.

Germination tubes were prepared by dispensing 7 ml of autoclaved 1% water agar into 15 ml conical tubes in a biosafety cabinet, avoiding air bubbles. A single sterilized seed was transferred to each tube with sterile forceps and overlaid with 40 µl of 10 mM MgCl₂. Tubes were sealed with AeraSeal membranes to permit gas exchange and held at 21.0°C in darkness for approximately 5 days, until the cotyledons had emerged and lifted clear of the agar surface. Seedlings were then transferred to a growth chamber held at 24.0°C with a 15 h light : 9 h dark photoperiod. Seedlings were 12 days old at first inoculation and remained in the same 1% water agar tubes, without added nutrients, for the duration of the experiment.

**Seedling inoculum preparation**

Bacteria were grown from frozen stock in 12 ml King's B medium in conical tubes at 28.0°C with shaking at ≥140 rpm for 24-48 h. Cultures were pelleted at 4,350 × g for 10–12 min, the supernatant discarded, and pellets resuspended in 10 mM MgCl₂, with the resuspension volume adjusted to culture turbidity so that low-density cultures were concentrated rather than diluted. Optical density was measured by transferring 100 µl of each suspension to a flat-bottomed 96-well plate alongside a 10 mM MgCl₂ blank, shaking for 5 s, and reading absorbance at 600 nm; blank values were subtracted. Suspensions were normalised to OD600 = 0.002 in two steps, first to OD600 = 0.02 and then by a 1:10 dilution, in 10 mM MgCl₂ containing 0.015% (v/v) Silwet L-77 (8 µl Silwet L-77 per 50 ml MgCl₂) to facilitate leaf colonization. Ten microliters of normalized suspension was applied directly to the seed leaves, with care taken to retain the droplet on the seed leaves. Control seedlings received 10 µl of 10 mM MgCl₂ with 0.015% Silwet L-77. All handling was performed in a biosafety cabinet; seedlings were allowed to dry before tubes were resealed with AeraSeal membranes and returned to the growth chamber.

*P. syringae* pv. tomato PT23 was used for seedling challenge rather than DC3000, which was used in the in vitro assays, to maintain comparability with the seedling and whole-plant protection assays previously established in this system [S16]. Both are *P. syringae* pv. Tomato, however, the strain PT23 has been demonstrated to confer repeatable diseased phenotypes in the seedling system. Inoculum density was fixed at OD600 = 0.002 throughout. Higher densities caused rapid necrosis in preliminary trials and were not used; this density reliably produces disease following *P. syringae* challenge while keeping total bacterial load well below that threshold.

**Mixed-model structure**

To test whether community filtrates differed from single-strain filtrates, spent-media class was fitted as a fixed effect with focal strain, plate nested within focal strain, and spent-media identity as random effects; spent-media identity was treated as random because community filtrates are largely drop-outs of a common parent community and are not independent. Because each community filtrate was assayed on a single plate, plate and community identity are partially confounded; we therefore tested for a systematic plate effect using fresh-medium controls run on both plates for every focal strain.

**Trait mixed-model set**

To test whether any measured trait explained variance in dAUC beyond strain identity, we fitted a pre-specified set of 20 mixed models sharing random intercepts for conditioner identity, recipient identity and plate, differing only in their fixed effects: standardized genomic traits (genome-predicted metabolic breadth, CAZyme repertoire, iron-acquisition gene content, genome size, GC content), biofilm formation, a composite growth-strategy axis, and nested taxonomic ranks, each entered on the conditioner side, the recipient side, or both, with an intercept-only model as the null. Models were fitted by maximum likelihood in lme4; variance components from the null model are reported as proportions of total variance.

**Seedling experiments: community panels and carrier solutions**

Experiment 1 tested five communities (Supplementary Table 6: AC, ABC, ABD, ACD, BCD). Experiment 2 tested the remaining ten communities (Supplementary Table 6: Full SynCom; A, B, C, D; AB, CD, AD, BC, BD). Experiment 3 compared four carrier solutions: Milli-Q water, Milli-Q water with Silwet L-77, 10 mM MgCl₂ with Silwet L-77, and 10 mM MgCl₂ with Milli-Q water and Silwet L-77.

**Choice of null expectations for community filtrates**

Because all cultures reached stationary phase in a common volume of medium, member contributions to resource depletion are not independent, and multiplicative (Bliss independence) expectations predicted near-complete growth suppression for the majority of communities; we therefore used the member mean and the most inhibitory member, which bracket the range of plausible expectations under shared resource limitation. Re-including the one excluded negative-control seedling (Experiment 2, day-0 score of 3) strengthens both principal seedling results (interaction P = 0.021; community-alone effect P = 0.005).

**Supplementary Results:**

This section gives detailed results summarized in the main text.

**Phylogenetic relationships and genome relatedness**

The phylogenetic relationships of the bacteria were highly congruent when using different sets of genetic data; however, the finer resolution at the tips of the tree varied. Phylogenetic relationships built using just 16S rRNA gene sequences place *P. asturiensis* SC10 and *P. rhodesiae* SC25 in a clade, however, when a set of 107 conserved single copy prokaryotic genes are used those two isolates are further diverged in the tree and *P. rhodesiae* clusters with *P. moraviensis* SC16 (Figure 1A, Supplementary Figure 1). Additionally, the relationships of the two *Pantoea* species varied, such that *P. allii* SC26 was positioned in a clade along with *E. tasmaniensis* SC04 when only the 16S gene is considered, however, when the conserved single copy gene alignment was used, the two *Pantoea* species, SC02 and SC26, clustered together. We calculated the pairwise whole genome average nucleotide identity (ANI) of all 17 members against one another (n = 136 comparisons), which revealed clustering patterns that reflected their phylogenetic history (Supplementary Figure 2). The ANI values revealed an average ANI = 67.6% across all comparisons with a maximum value of 98.1% (*P. agglomerans* SC02 comparison against *P. allii* SC26) to a minimum of 62.9% (*P. agglomerans* SC02 comparison against *E. sibiricum* SC13).

**Shared and strain-specific gene content**

Because our aim was to test whether interaction outcomes are predictable from genome content, we first asked how much functional content these strains actually share. We identified KEGG pathways in our community using BlastKOALA which successfully annotated an average of 53.7% of genes in each genome (ranging from 39.7% to 67.3% annotated across all genomes). Phylogenetic orthology inference recovered 495 orthogroups present in all sixteen commensals and 432 across all seventeen genomes including the pathogen, roughly a tenth of the approximately 4,400 open reading frames in an average genome of this panel. The great majority of gene content is therefore lineage- or strain-specific, which is why we characterize strain-level capacity below rather than treating the community as functionally uniform, and it sets the expectation that any predictive signal should come from where strains differ rather than from what they share.

**Predicted amino acid biosynthesis and carbon catabolism**

We identified amino acid biosynthesis in all the strains, revealing 9 out of 17 strains with high confidence pathways across all of the amino acid pathways. The isolate *E. sibiricum* (SC13) was the only strain found to have medium to low confidence for more than half of the amino acid pathways (6 = low confidence and 6 = medium confidence). Clustering by principal components analysis (PCA) revealed similar amino acid biosynthesis metabolic profiles among the Gram-negative strains while the Gram-positive strains had high variability across PC1 and PC2 (Supplementary Figure 3). We also predicted the ability of the strains to catabolize small carbon sources which revealed more metabolic diversity across the strains compared to amino acid biosynthesis. For small carbon source utilization, the Gram-negative strains displayed higher variability along PC1 where the Gram-positive bacteria had more similar predicted metabolic potential (Supplementary Figure 4).

**Carbohydrate-active enzymes and iron-related genes**

As phyllosphere bacteria encounter polysaccharides in their natural environment like cellulose, we annotated and quantified the carbohydrate-active enzymes (CAZymes) in each of the genomes. We identified a range of 55 (*E. sibiricum* SC13) to 147 (*R. festucae* SC21) CAZymes, with an average of 90 CAZymes per genome (Figure 1A). The number of CAZymes in a single genome was not correlated with genome size (R = -0.21, P = 0.41). We also interrogated the genomes for genes related to iron acquisition, oxidation/reduction, and storage. Iron competition is one of the few antagonistic mechanisms that operates without cell contact, and the siderophores that mediate it persist in cell-free filtrates, so iron-acquisition capacity is a candidate predictor of the interactions measured here. Isolates that were closely related based on phylogenetics shared similar numbers of iron-related genes, with the *Pseudomonas* genomes harboring the highest number, an average of 129 genes compared to 61 for all others (Figure 1A). These counts index annotated capacity rather than realized function: about half the coding capacity of these genomes is unannotated, and carriage of iron-acquisition genes does not imply siderophore production under any given condition.

**Secretion systems and biosynthetic gene clusters**

Finally, we interrogated the genomes for evidence of secretion systems, which have many roles relevant to nutrient depletion including resource scavenging [S17], as well as for evidence of secondary metabolite biosynthetic gene clusters (SM BGCs) that could be relevant to interaction outcomes. We identified several secretion systems across the genomes with most occurring in the Gram-negative species, likely due to membrane biology and database limitations (Supplementary Figure 5). The most abundant system was the T5aSS, which was also the most abundant secretion system found in a dataset of over 1,000 bacterial genomes [S18], followed by the T1SS and T4ap. A total of 138 SM BCGs was identified amongst the 17 genomes, with an average of 8 per genome, a maximum of 13 (*B. thuringiensis* SC35), and a minimum of 3 (*E. sibiricum* SC13). Using a similarity cutoff of 50% left 31 high confidence SM BCGs.

**Growth phenotypes**

Specifically, we measured growth rate, biofilm formation, and carbon-source utilization across all strains; these traits are later used to interpret species-specific contributions to community dynamics and functional outputs. Using a medium that supports the growth of all the strains, King’s Broth media, we measured the growth rates of each bacterium over 72 hours (Supplementary Figure 6). All strains reached stationary phase within the monitored time period, however, the strains displayed diverse growth characteristics (Table 1).

**Plate diagnostics and variance partitioning in the spent-media assay**

A plate diagnostic comparing each focal strain’s mean AUC on its two assay plates found no systematic plate effect (median difference = 1.6%, paired Wilcoxon P = 0.49, Supplementary Figure 10), so plate confounding does not explain this difference. Of the total variance in dAUC, spent-media identity accounted for 18.2% and focal-strain identity for 21.3%, and the remaining 58.9% was residual well-to-well variation. Variation among community treatments is therefore attributable to which filtrate was applied and which focal strain was tested, rather than to plate-level technical variation.

**Trait associations: statistical treatment and the metabolic breadth measure**

We report these associations as effect sizes rather than as hypothesis tests, and we do not apply a multiple-comparison correction across the panel. The twenty comparisons are not twenty independent questions: the two axes are two views of a single strain-level property, and the traits themselves have some associations such as genome size, iron-acquisition gene content and metabolic breadth rise in step, and total growth and maximum density are close to the same measurement (Figure 1B). With seventeen strains the panel is in any case better read as a description of the direction and relative magnitude of trait associations than as a set of tests, and we claim no individual association as significant. Whether any trait explains variance beyond strain identity is a separate question, addressed below at the level of individual wells. Two features of the metabolic breadth measure limit its interpretation: it is scored over only the 13 EcoPlate substrates for which a genomic prediction was available, and four strains are predicted to use 12 of these 13, so the measure has little power to separate the most metabolically versatile strains. It is also collinear with Gram status across this panel (ρ = +0.83), which is unsurprising given that the Gram-negative members are predominantly copiotrophic Pseudomonadota. Breadth nonetheless explains sensitivity better than Gram status alone (ρ = +0.56 against +0.43), and the association holds within the Gram-positive group but is weaker within the Gram-negative group (ρ = +0.61, n = 8 and ρ = +0.43, n = 9), so we report it as a metabolic association while noting that a 17-member panel cannot separate metabolic capacity from broad taxonomy.

We treated the two directions separately because the interaction matrix is substantially asymmetric (mean |S - Sᵀ| = 0.30 against a total range of 1.50), so collapsing it into a single symmetric distance would discard one direction of every pairwise interaction.

**pH and suppression of *P. syringae***

Filtrates that were indistinguishable in pH nonetheless differed widely in suppression: among the eleven filtrates measured at pH 6.5, *P. syringae* dAUC ranged from -0.06 to -0.97, and among the ten measured at pH 6.33 it ranged from +0.14 to -0.76.

**Reproducibility of community-mediated pathogen suppression**

To assess reproducibility, we repeated the assay with newly generated biological replicates of 22 community filtrates (n = 66; Methods), whose composition was profiled by 16S rRNA gene sequencing (Supplementary Figure 14). We found a significant difference in final community composition among the different built consortia (R2 = 0.70, P = 0.001, Adonis PERMANOVA; Supplementary Figure 15), with a non-significant result for the dispersion test (P = 0.979) supporting biologically driven outcomes not due to heterogenous variability or noise. The alpha-diversity, measured with the Shannon Index, was also significantly different amongst the different built consortia (P = 3.01 × 10⁻⁵, ANOVA) with lower alpha-diversity for the groups that excluded more members: for example, the communities that excluded all Gram-negative strains had the lowest alpha-diversity (Supplementary Figure 16A). Evenness differed significantly among constructed consortia (P = 0.0036, ANOVA) and increased with the number of strains present (P = 1.7 × 10⁻⁵, linear regression), indicating that larger consortia supported more even coexistence among persisting strains (Supplementary Figure 16B).

Across the four focal strains tested (Methods), the replicate filtrates produced an average dAUC = -0.15 (Supplementary Figure 17). To evaluate reproducibility of relative growth change across experiments, we compared dAUC values from the replicate spent-media assay to those obtained in our initial experiment. Across all focal strains and community filtrates, relative growth change values were significantly correlated (Spearman ρ = 0.21, P = 6.3 × 10⁻⁴), indicating consistent relative suppression across independent batches of spent media. The mean dAUC varied modestly among community spent-media groups, ranging from -0.26 (Full-SC02) to -0.06 (Full-SC13). The most suppressive filtrates in this replicate were Full-SC02, Full-SC04, and Full-SC03, followed by Full-SC25, Full-SC26, and the full synthetic community filtrate (Full), whereas Full-SC13 was the least suppressive on average (Supplementary Figure 18).

**Additional seedling control arms**

Challenging twice raised them further (Experiment 1, +1.48; Experiment 2, +1.00), and a challenge delivered at day 0 rather than day 3 also produced elevated scores (+1.02), consistent with younger seedlings being more susceptible.

**Community pre-inoculation and seedling symptoms without the pathogen**

Pooled across all 15 communities, seedlings pre-inoculated with a community but not challenged with the pathogen scored modestly higher than untreated seedlings (+0.92 versus +0.59 baseline-adjusted, P = 0.010; main-text Fig 5B), as might be expected of strains isolated from mature plants rather than selected for seedling compatibility. Four communities, including the full PhylloStart community, reached nominal significance individually (P = 0.023–0.048), but none survived correction for multiple testing, so we report this effect only at the panel level. It was not due to the inoculation carrier: four carrier solutions, including the MgCl₂ and Silwet L-77 mixture used throughout, were indistinguishable from one another and from untreated seedlings (Kruskal–Wallis P = 0.49; Supplementary Figure 19). Because a community alone raises scores toward those of the pathogen alone, the smaller pathogen effect in pre-colonized seedlings may reflect restricted pathogen damage or a ceiling on symptom expression; scores stayed well below the scale maximum, but quantifying pathogen load will be needed to distinguish these.

**Phenotypes of the strongest single-strain suppressors**

We next asked whether the two strongest single-strain inhibitors (SC13 and SC35) shared common phenotypic traits. Inhibitory strength was not associated with doubling time (Wilcoxon W = 7, p = 0.29), carbon-use breadth (W = 16.5, p = 0.88), or biofilm formation (Fisher’s exact test p = 0.51), indicating that strong single-strain suppression is not explained by growth strategy or metabolic breadth.

**Iron-acquisition capacity and conditioning**

The expectation that greater iron-acquisition capacity would confer stronger conditioning of a shared medium is not supported: the association runs to sensitivity rather than conditioning, meaning that strains carrying more iron-acquisition machinery were less affected by depleted media rather than more depleting of it. The same holds at the level of gene counts: the four *Pseudomonas* commensals carry roughly twice the iron-related gene content of the rest of the panel (129 against 61 on average) yet show conditioning effects that overlap the remainder of the panel.

**Pathogen growth in drop-out filtrates retaining SC13**

In the filtrates lacking SC10, SC14 or SC16, *P. syringae* grew at least as well as in fresh medium despite *E. sibiricum* SC13 being present.

**Supplementary tables:**

See excel file “2026-11-10_SupplementaryTable_File.xlsx”

**Supplementary figures:**

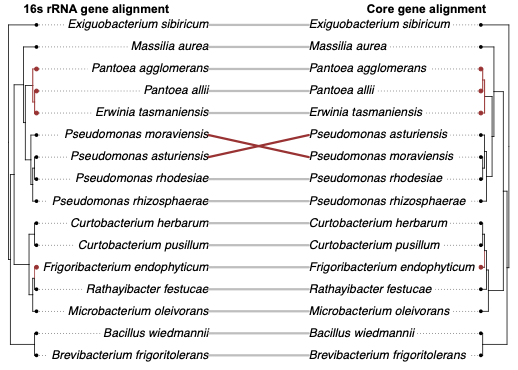

**Supplementary Figure 1:** **Comparison between two phylogenetic trees depicting the relationships between the 16 phyllosphere bacteria that make up PhylloStart.** The tree on the left was built using the 16s rRNA gene alignment (left) and the tree on the right was built using a core gene alignment of roughly 100 highly conserved prokaryotic genes.

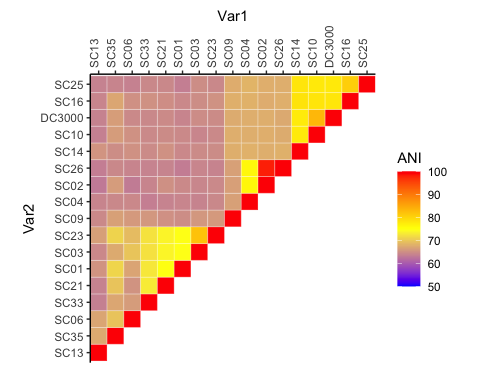

**Supplementary Figure 2:** **Pairwise average nucleotide identity (ANI) for all 17 bacterial species analyzed in this paper.** Warmer colors represent higher similarity and cooler colors represent less shared identity.

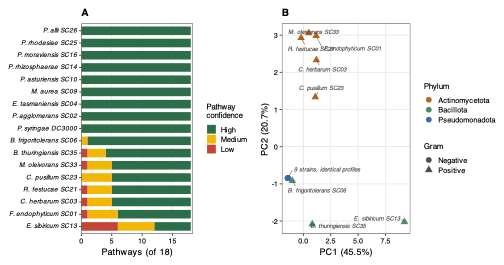

**Supplementary Figure 3:** **Predicted amino acid biosynthesis capability across the 17 isolates.** (A) Number of amino acid biosynthesis pathways recovered per strain, coloured by GapMind confidence call (high, medium, low). (B) Principal components analysis of the confidence-scored pathway profiles, with points coloured by phylum and shaped by Gram status; axis labels give the percentage of variance explained by each component. Nine of the 17 strains returned high-confidence calls for every pathway, and *E. sibiricum* SC13 was the only strain with medium- or low-confidence calls for more than half of them.

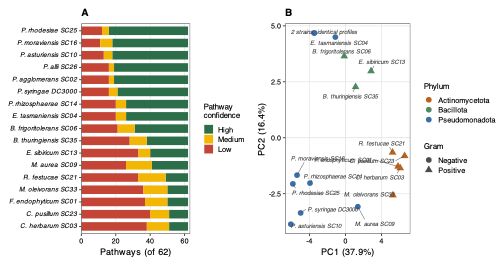

**Supplementary Figure 4:** **Predicted small carbon source utilization across the 17 isolates.** (A) Number of catabolic pathways recovered per strain, coloured by GapMind confidence call (high, medium, low). (B) Principal components analysis of the confidence-scored pathway profiles, with points coloured by phylum and shaped by Gram status. Predicted small carbon utilization was more variable across strains than amino acid biosynthesis, with the Gram-negative isolates spreading along PC1 and the Gram-positive isolates occupying a narrower region of the ordination.

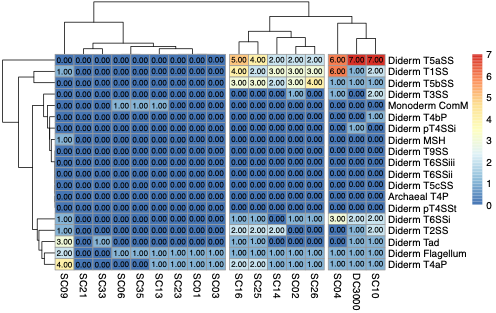

**Supplementary Figure 5**: **Heatmap depicting the number of secretion systems annotated by TXSS for the 17 bacteria species.**

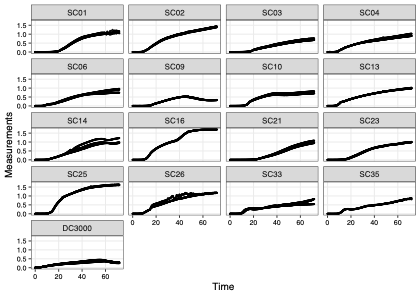

**Supplementary Figure 6:** **Growth curve dynamics of all the 17 strains used in the spent media analyses.** Growth curves were performed in King’s B liquid media in 96-well plates with optical density (OD600) measurements taken every 15 minutes for 72 hours at 28.0°C.

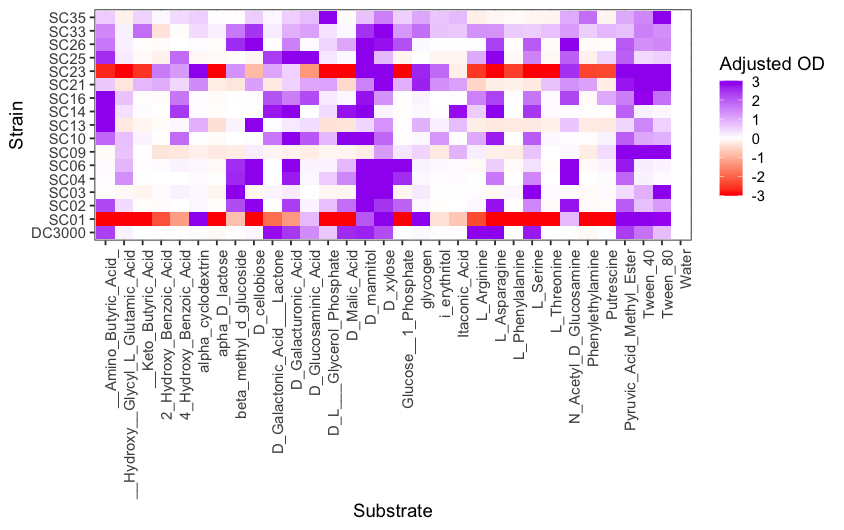

**Supplementary Figure 7:** **Metabolic profiling using Biolog EcoPlates.** Carbon substrate utilization profiles for each strain across 31 carbon sources. The heatmap depicts the relative color development adjusted OD values for each well after incubation. Distinct metabolic fingerprints highlight strain-level functional diversity within the SynCom.

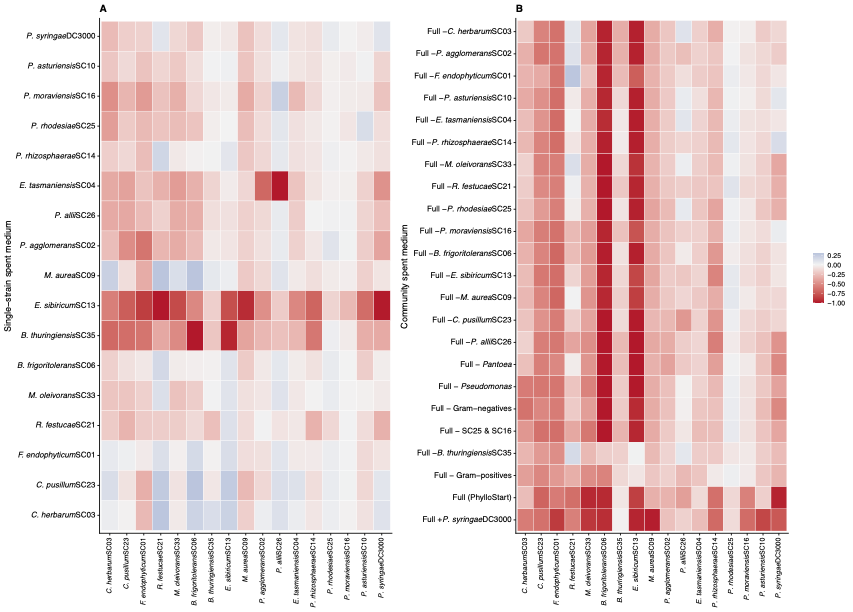

**Supplementary Figure 8: Distribution of relative growth change (dAUC) across all spent media and focal strain combinations.** dAUC is growth in a spent medium relative to that strain's growth in fresh King's B medium, calculated as (spent − fresh) / fresh, so zero indicates growth matching fresh medium and negative values indicate suppression. The mean across all combinations was dAUC = −0.29.

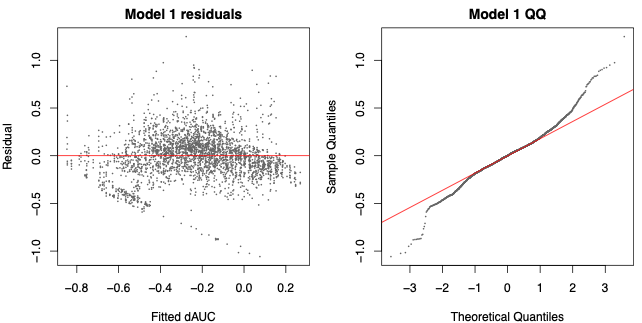

**Supplementary Figure 9: Diagnostic plots for the linear mixed-effects model of dAUC.** Residual and quantile plots for the model fitting media class (community versus single-strain) as a fixed effect, with focal-strain identity, plate nested within focal strain, and spent-media identity as random effects. Under this model, community filtrates reduced growth by an average of 0.203 dAUC units more than single-strain filtrates (df = 38.5, P = 2.2 × 10⁻⁵).

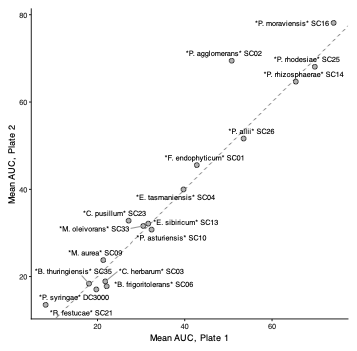

**Supplementary Figure 10: Plate diagnostic based on fresh-medium controls.** Each focal strain was assayed on two 96-well plates, with fresh King's B medium run on both. Mean fresh-medium AUC is compared between the two plates for each of the 17 focal strains. The median difference was 1.6% (paired Wilcoxon P = 0.49), indicating no systematic offset between plates and therefore no plate confounding of the community filtrate comparisons.

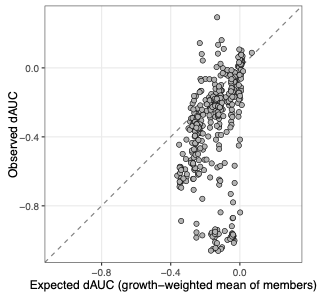

**Supplementary Figure 11: Observed dAUC against a growth-weighted averaging null.** For each of the 391 community by focal strain combinations, the observed dAUC is plotted against the expectation obtained by averaging the member dAUC values for that community, weighting each member by its own AUC in fresh King's B medium. The dashed line is the 1:1 expectation; points below it indicate suppression stronger than expected from the members in isolation (Wilcoxon signed-rank P = 2.9 × 10⁻⁴²). Weighting members by their growth does not change the conclusion drawn from the unweighted averaging null in Figure 3A.

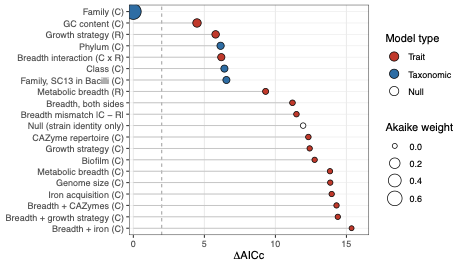

**Supplementary Figure 12: Conditional R² across the candidate set of mixed models.** Twenty models sharing random effects for conditioner identity, recipient identity and plate were fitted with different combinations of genomic and phenotypic traits as fixed effects. Conditional R² was 0.432 to 0.436 for every model, including the intercept-only null, indicating that trait terms redistribute variance already attributable to conditioner identity rather than explaining variance beyond it. Model specifications and AICc values are given in Supplementary Table 5.

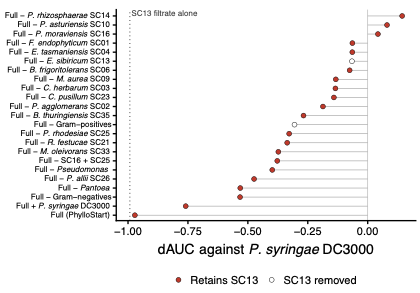

**Supplementary Figure 13:** **Community effects on *P. syringae* DC3000 shaded by whether the community retains *E. sibiricum* SC13.** The dotted line marks the effect of the SC13 filtrate alone on the pathogen.

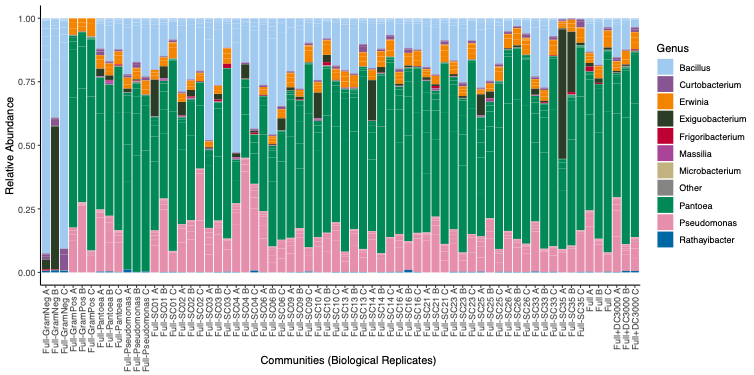

**Supplementary Figure 14:** **Relative abundances are shown for each of the three biological replicates per community type.** Each bar represents the ASV composition (assigned via DADA2 and SILVA) immediately prior to preparing spent media. Communities differed significantly by treatment (PERMANOVA R² = 0.70, P = 0.001), while dispersion did not differ (P = 0.979), indicating biologically meaningful differences in composition rather than variability in replicate spread. *Pantoea agglomerans* SC02, *Pantoea allii* SC26 and *Erwinia tasmaniensis* SC04 (family Erwiniaceae) are identical or nearly identical across the V3–V4 region and cannot be reliably distinguished from one another in these profiles.

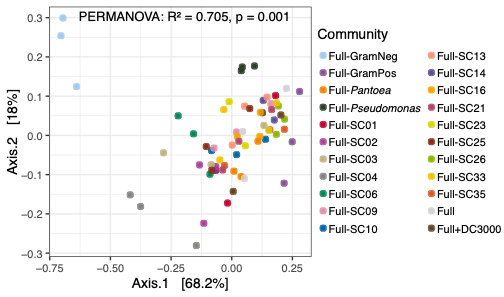

**Supplementary Figure 15: Principal coordinate analysis based on Bray–Curtis dissimilarities of the community spent medias.**

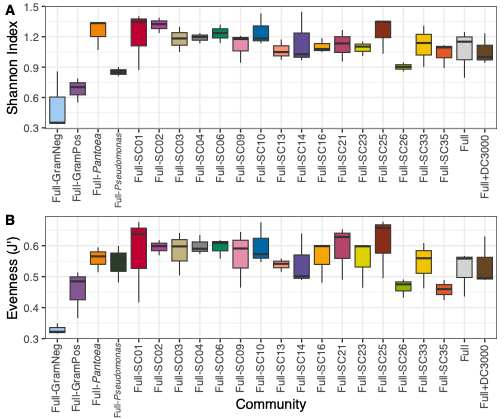

**Supplementary Figure 16.** **Alpha diversity of the biologically replicated community cultures.** A) Shannon diversity and B) Pielou’s Evenness (J’) of bacterial communities across community spent-media types prior to filtration.

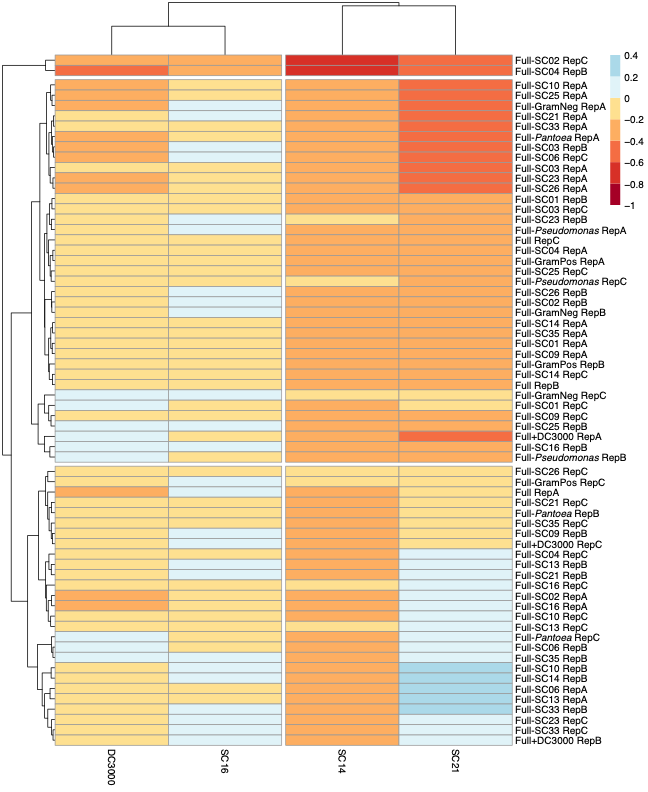

**Supplementary Figure 17:** **Heatmap depicting the relative growth change for all biologically replicated community spent media (rows) that were reconstructed and retested against 4 focal species (columns).** For each community, three biological replicates were made (see Methods) and are depicted as RepA-C to represent the three replicates. Warmer colors depict stronger relative growth change (more negative interactions) and cooler colors depict neutral to more positive interactions. Unlike the primary experiment, these filtrates are biological replicates: each community was conditioned in three independently grown cultures, giving 66 independently generated community spent media.

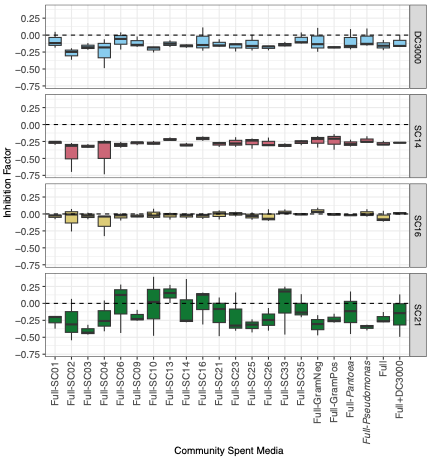

**Supplementary Figure 18: Boxplots depicting the biological replication of community spent media effects and reveals consistent inhibition patterns across focal strains.** These are the biologically replicated filtrates described above, comprising three independently conditioned batches of each community, and are distinct from the single conditioning batch used in the primary experiment.

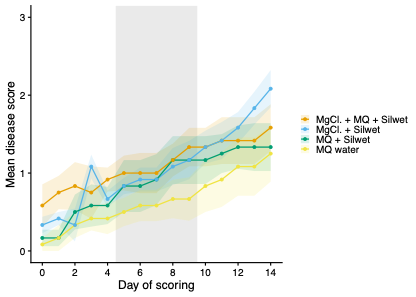

**Supplementary Figure 19: Seedling disease scores are unaffected by the inoculation carrier solution.** Baseline-adjusted symptom scores for seedlings treated with each of four carrier solutions, including the MgCl₂ and Silwet L-77 mixture used throughout this study, alongside untreated seedlings. The four carriers were indistinguishable from one another and from untreated seedlings (Kruskal–Wallis P = 0.49).
